# Mitotyping – An integrative framework to quantify mitochondrial specialization and plasticity

**DOI:** 10.1101/2025.02.03.635951

**Authors:** Anna S. Monzel, Jack Devine, Darshana Kapri, Gabriel Sturm, Caroline Trumpff, Vilas Menon, Philip L. De Jager, Jose Antonio Enriquez, Martin Picard

## Abstract

Mitochondria are a diverse family of organelles with highly specialized functions. While they share common features, their molecular and functional diversity remains underexplored. Here, we introduce a quantitative pipeline to define the degree of molecular specialization among different mitochondrial phenotypes, or *mitotypes*. By distilling hundreds of validated mitochondrial genes into 149 biologically interpretable MitoPathway scores, this streamlined mitotyping framework enables investigators to quantify and interpret mitochondrial diversity and plasticity from transcriptomic data across a variety of natural and experimental contexts. Using this approach, we show that mouse and human multi-organ mitotypes segregate along two main axes of mitochondrial specialization, characterize robust longitudinal and perturbation-induced metabolic plasticity in cultured human fibroblasts, and resolve cell-type and single-cell mitochondrial recalibrations across the human brain and sperm developmental trajectory. Together, this framework provides a practical and extensible approach for analyzing mitochondrial specialization in complex biological systems.

## Introduction

Much of a cell’s metabolic activity centers around the mitochondrion – a metabolic hub and signaling platform integrating numerous cellular and organismal behaviors^1,2^. Like all cell (sub)types, mitochondria arise from a single common ancestor, the “mother” mitochondria of the oocyte (**Figure 1A**). The mitochondrial phenotype of human oocytes is rather unique, suppressing both complex I gene and protein expression^3^. As organs and tissues develop and cell type identities are established, mitochondria likewise adapt and evolve with the cell, dynamically meeting its changing metabolic demands^4–6^. These adaptations give rise to the full spectrum of terminally differentiated mitochondrial phenotypes identified to date, reflecting a form of horizontal developmental plasticity. Different mitochondrial phenotypes are even observed among different sub-compartments of the same cell ^7–10^. Temporally, it is also established that exercise^11,12^, nutritional interventions^13^, hormones^14^, and subjective experiences including psychosocial factors^15–17^ are linked to specific functional recalibrations in mitochondrial biology, demonstrating how plastic mitochondria are.

**Figure 1:**
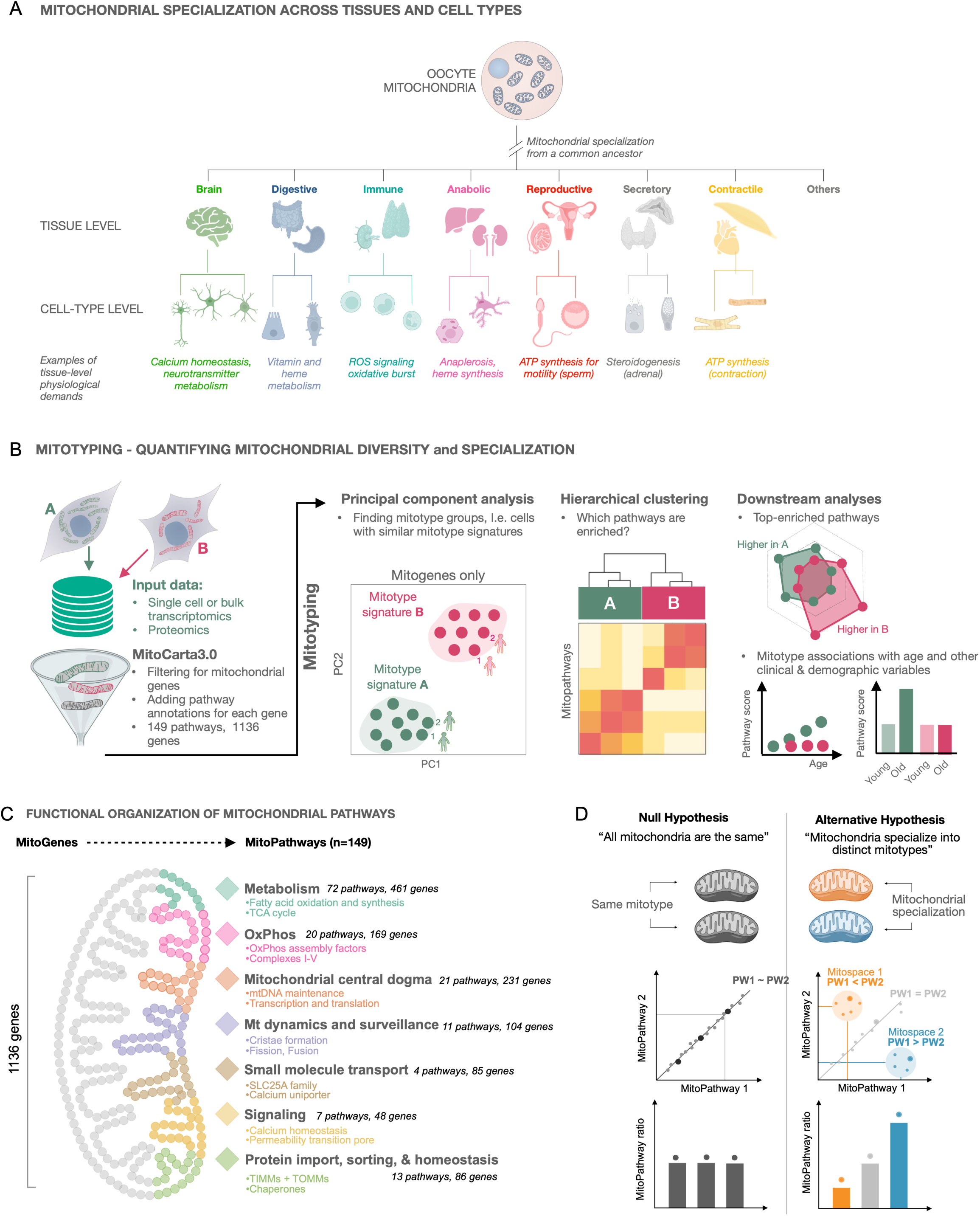
Quantifying mitochondrial specialization. (**A**) Mitochondrial specialization can be quantified at different levels of complexity. All mitochondria originate from the maternal mitochondrial population in the oocyte. During development, mitochondria must acquire tissue-specificity. Tissue-level mitotypes reflect mitochondrial specialization across the mammalian body and are a combination of multiple cell type-level mitotypes. Cells rely on specialized mitochondria that house a variety of pathways vital for cellular function. (**B**) Mitotyping quantifies mitochondrial specialization from omics data. Mitochondrial genes or proteins are extracted from normalized transcriptomics or proteomics data, and MitoPathway annotations are added using MitoCarta3.0^20^. In each tissue, cell type or cell state, the mitochondrial transcriptome or proteome is deeply characterized using MitoGenes or MitoPathways to find distinct mitotype signatures, identify groups with similar mitotypes or to study mitochondrial adaptation. **C)** Mammalian mitochondria encompass >1100 genes and proteins, the majority of which are encoded in the nuclear genome and a small fraction (13 genes) in the mitochondrial genome. The expression of MitoGenes and MitoProteins varies in a context-dependent manner depending on the environment. Different sets of MitoGenes and MitoProteins form MitoPathways that are enriched in different tissues, cell types, or cell states. We use MitoPathway scores based on MitoCarta3.0 ^20^annotations to quantify mitotype differences. **D)** Conceptual model of mitochondrial specialization on the molecular level. *Left panel*: Under a uniform model, all MitoPathways scale proportionally with mitochondrial content, and ratios of two MitoPathway scores would be the same. *Right panel*: Under mitochondrial specialization, specific pathways are selectively up- or downregulated, positioning cells or tissues within distinct regions or *mitospaces*. Ratios of a set of two MitoPathways differ between the groups.

Most studies pragmatically assess mitochondrial biology using a limited set of molecular features, activities, or functional parameters. Such a focus can be misleading as a change in one or two parameters in isolation can appear dysfunctional, whereas a broader view can instead reveal adaptive recalibrations^18^. For instance, a decline in one function alongside many others may reflect a decline in mitochondrial mass; but if that same function declines while others are selectively upregulated, it may reflect a purposeful recalibration. Most techniques often fail to distinguish between these scenarios, conflating *adaptive recalibration* with *dysfunction*^19^.

To address this limitation, we propose a systems-level framework that integrates all known mitochondrial pathways, and identifies which ones are prioritized in a given context. We term this approach *mitotyping*. Mitotyping extracts pathway-level information from omics data to molecularly phenotype mitochondria (**Figure 1B**). It works by computing MitoPathway scores based on the expression of genes within the 149 curated mitochondrial pathways defined by MitoCarta3.0^20^, each reflecting specific mitotype features, functions, and behaviors (**Figure 1C**). By contrasting these scores between one another, we then quantify how specific mitochondrial functions are expressed and prioritized in each sample. Under the null hypothesis where all mitochondria are uniformly specialized, cells or tissues with more mitochondria would show proportional increases in all pathway scores (**Figure 1D, *left*).** In contrast, if mitochondria are functionally specialized, certain pathways will be upregulated relative to others, shifting samples into distinct regions of the possible mitotype space (**Figure 1D, *right***).

Here, we show how mitotyping of transcriptomic datasets across multi-tissues and single cells captures mitochondrial and related recalibrations across the mammalian body, in distinct cell states and developmental trajectories, and in response to experimental perturbations *in vitro*. By identifying pathway-level prioritization, mitotyping quantifies stable mitochondrial signatures, transient states, and patterns of mitochondrial plasticity that remain inaccessible with conventional analyses.

### Organ and tissue-based mitochondrial profiling

#### Mouse tissue-specific mitotypes

Using the original MitoCarta mouse proteome dataset^6^, we analyzed mitochondrial protein abundance (n=977 proteins) across 14 well-defined tissues (**Figure S1A**). Hierarchical clustering of mitochondrial genes alone – *without* any canonical tissue-specific or cell-type specific marker – yielded clusters of tissues that matched broad functional categories such as brain, contractile muscles, anabolic tissues, and all other tissues, demonstrating that mitochondrial features alone were sufficient to recover biologically meaningful groups (**Figure 2A**). A principal component analysis (PCA) illustrated particularly clearly how tissues from the brain exist as a distinct mitotype cluster, segregating away from the anabolic liver and kidney, and from the contractile mitotype signature of cardiac and skeletal muscles. A fourth, more heterogeneous group included digestive, reproductive, and adipose tissue mitotypes (**Figure 2B**). By contrast, variation along PC3 and PC4 did not reveal additional coherent axes of separation, suggesting that the dominant structure of mitotype diversity across tissues is effectively captured by the first two principal components (**Figure S2A**).

**Figure 2:**
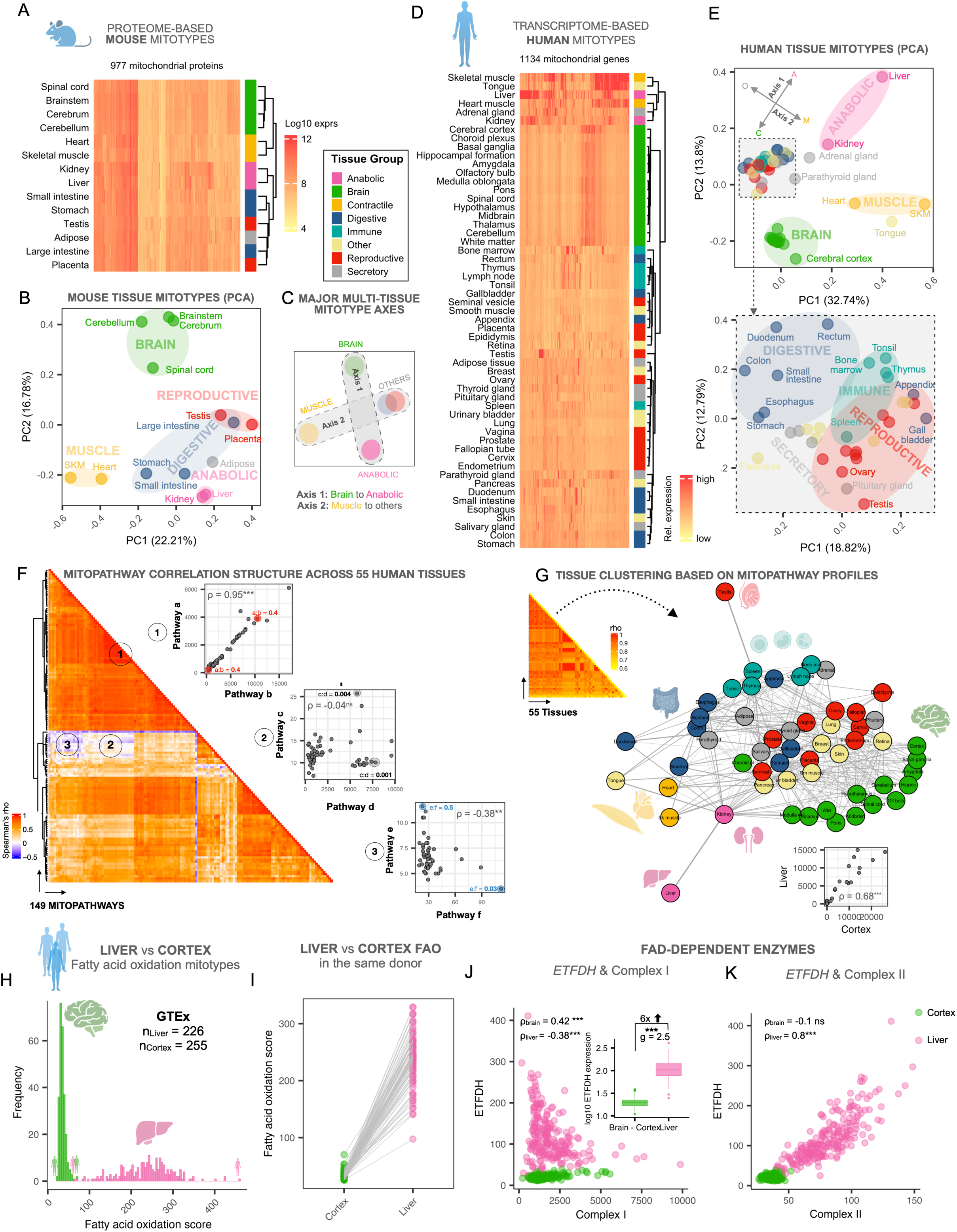
Mouse and human tissues contain molecularly distinct molecular mitochondrial phenotypes. **A)** Hierarchical clustering and **B)** Principal component analysis (PCA) of the 14 mouse mitochondrial proteomes from MitoCarta3.0 ^20^reveals the grouping of physiologically-related tissues, based uniquely on mitochondrial proteins. **C)** Major multi-tissue mitotype axes as suggested by the PCA in B. **D)** Human mitotypes based on consensus gene expression data from the human protein atlas (HPA) and the Genotype-Tissue Expression (GTEx) project ^21,23,70,71^, mapped to the human MitoCarta3.0 gene list. Relative expression for each mitochondrial gene (columns) in 55 human tissues (rows), color-coded by physiological systems. **E)** PCA of all human mitotypes (top) and of the tissue subsets highlighted in the box (bottom), illustrating the mitotype-based clustering of tissues with similar physiological functions. The PCA shows the same major tissue axes as in C (arrows in top panel). Axis 1 separates the CNS from anabolic tissues, and axis 2 separates muscle from other tissues. **F)** Spearman’s ρ correlation-based MitoPathway analysis shows that not all MitoPathways are tightly correlated across 55 human tissues from the Human Protein Atlas dataset. On the right side of the heatmap are representative bivariate MitoPathway plots of 1) positively correlated, 2) not correlated, and 3) negatively correlated MitoPathways. In addition, numbers indicate MitoPathway ratios for some example tissues that occupy different mitospaces. **G)** Adjacency matrix presentation of between-tissue spearman correlations across 55 tissues and 149 MitoPathways demonstrating similar group clusters as on the gene level. The bivariate plot on the bottom shows a representative association between liver and cerebral cortex. **H)** Fatty acid oxidation (FAO) in tissues from axis 1 (cerebral cortex vs liver) across >220 human donors from the GTEx project showing higher expression of FAO in the liver relative to the cortex. **I)** FAO in donor-matching samples from cortex and liver showing that liver FAO score is on average 7.1x higher than in the cortex **J)** Bivariate plot of FADH-dependent electron transfer flavoprotein dehydrogenase *ETFDH* and Complex I (CI) expression score and boxplot of *ETFDH*/CI ratios in cerebral cortex vs liver. The liver expresses 6x more ETFDH compared to the brain. **K)** Bivariate plot of *ETFDH* and Complex II (CII) expression score in cerebral cortex vs liver. The liver shows a tight correlation between FADH-dependent enzymes and CII expression. Abbreviations: *SKM*, skeletal muscle; Hierarchical clustering methods in A and D using the Euclidean distance and the Ward’s D2 hierarchical Agglomerative Clustering Method^72^. Brunner Munzel test and Hedges’ g for effect size. \*\**p*<0.01, \*\*\**p*<0.001.

This integrative analysis pointed to two main conclusions: First, mitochondrial proteins alone are sufficient to distinguish tissue types, revealing the robustness of the mitotype proteomic signatures. Second, the cardinal alignment of tissues along four poles suggested the presence of *two major axes* (brain, contractile muscles, anabolic tissues, and all others) of mitochondrial specialization (**Figure 2C**).

#### Human tissue-specific mitotypes

We repeated this analysis using mitochondrial RNA expression across 55 human tissues from the Human Protein Atlas (HPA)^21,22^ and Genome-Tissue Expression (GTEx)^23^ consensus databases (**Figure S1B**). As in mice, the unsupervised analysis recapitulated the clustering of brain, anabolic, and contractile tissues, and a mixed group of the digestive, reproductive, endocrine, and immune system tissues (**Figure 2D**). Here also, tissues aligned along the same two axes of mitochondrial specialization extended from the most anabolic of all organs, the liver, to the most catabolic, the brain (*Axis 1*); and from contractile muscle tissues (heart, skeletal muscle, tongue) to other tissues (*Axis 2*) (**Figure 2E** and **S2B**). Further analysis of “other” tissues by PCA showed remarkably coherent mitochondrial transcriptional signatures for most digestive tube segments, for most immune tissues (except the spleen), and for reproductive or secretory tissues. Thus, the linear combination of mitochondrial genes alone is sufficient to distinguish major tissue types.

To better understand the molecular basis of these mitotype differences, we next examined how individual mitochondria pathways, as annotated in MitoCarta3.0^20^, are correlated across tissues. We found a core set of “*canonical*” MitoPathways that were tightly correlated, i.e. tissues with more mitochondria have proportionately more of these pathways (*Example 1* **Figure 1F**). In contrast, other pathway combinations showed weak or no correlation (*Example 2*), and some were even negatively correlated (*Example 3*). This suggests potential biological tradeoffs or functions carried out by specialized mitochondria in specific contexts. Taken together, the clustering of tissues based on mitochondrial genes alone above (Figure 2A-E) and the pathway-based correlations collectively refute the null hypothesis (“all mitochondria are equal”) which would predict that tissues cluster randomly.

To pressure test this idea, we extended the multivariate analysis at the pathway level using tissue-to-tissue correlations with the 149 MitoCarta pathways as sole input. The network structure of all 55 tissue correlations illustrated distinct clusters of brain, anabolic, contractile, immune, and a mixed group of reproductive, secretory and other tissues (**Figure 2G**)—similar to the gene-level grouping. This aligns with previous work^5,24–26^ (also reviewed in ^4,19^), pointing at the existence of somewhat different human mitochondrial phenotypes, or families of organelles, which can be mapped by the combinations of the 149 interpretable MitoPathways.

#### Pathway level mitotyping

To grasp the magnitude of the molecular specialization between tissues, we examined the expression of the well-characterized MitoPathway fatty acid oxidation (FAO) in GTEx (n=226-255), comparing the brain known to have low-to-absent FAO capacity^27^ to the liver that readily oxidizes lipids for oxidative phosphorylation (OxPhos) and anaplerosis^28,29^ (**Figure S1C**, **S2C-D, S3** and **2H)**. The mean difference in fatty acid oxidation score between these tissues was 6.7-fold (*p*<0.0001, **Figure S2E**). Comparing FAO expression in the liver and cortex from the same donor, we showed that as expected from the population-level data, the FAO score was *always* ∼3-11x higher in the liver than in the brain (average=7.1-fold, **Figure 2I**), establishing the reproducibility and magnitude of differences in mitochondrial gene expression between these human tissues. This result illustrates both the quantitative differences between tissues, and the internal consistency (in 100% of donors, n = 79) of mitotype signatures across individuals.

#### Gene-pathway level mitotyping

To further validate tissue-specific differences in alternative electron-entry routes into the electron transport chain (ETC), we mapped the FAO-ETC relationships in liver and brain at both the gene and pathway levels. FAO delivers electrons to the ETC by oxidizing acyl-CoAs in the mitochondrial matrix through the TCA cycle and via the FADH-dependent electron transfer flavoprotein dehydrogenase ETFDH. ETFDH transfers electrons directly to Coenzyme Q (CoQ), bypassing Complex I (CI)^30,31^. Consequently, when FAO activity is high, a smaller proportion of total electrons enter CI. Accordingly, we expected liver mitochondria – where FAO is robust – to express more *ETFDH* relative to CI, while maintaining proportional expression with Complex II (CII). Consistent with this prediction, in liver tissues *ETFDH* expression was negatively correlated with CI (ρ = - 0.38, *p*<0.001, **Figure 2J**) and strongly positively correlated with CII (ρ = 0.8, *p*<0.001, **Figure 2K**). In contrast, in the brain, where FAO is limited, *ETFDH* expression was very low (∼6-fold lower than liver) correlated positively with CI (ρ = 0.42, *p*<0.001, Figure 2J), and showed no correlation with CII (Figure 2K), which is also expressed at low levels in brain mitochondria. Together, these patterns highlight a quantifiable tissue-specific coordination between FAO capacity and alternative electron-entry routes into the ETC at the gene-pathway level, consistent with distinct electron-flow priorities in liver vs brain mitochondria.

#### Gene level mitotyping

Building on our pathway- and gene-pathway-level mitotyping analyses, we next focused on gene-level mitotyping to specifically examine enzymes that inject electrons into the ETC while bypassing CI, including CII (SDHA-D), dihydroorotate dehydrogenase (DHODH), choline dehydrogenase (CHDH), prolyl dehydrogenase (PRODH), hydroxyproline dehydrogenase (PRODH2), and sulfide quinone oxidoreductase (SQRDL). We showed that FAD-dependent enzymes are expressed at lower levels in brain compared to liver mitotypes (**Figure S2F**) with PRODH as the only exception, likely reflecting its role in glutamate synthesis^32^. This observation prompted us to further evaluate mitotype adaptation to the branched electron entry points of the ETC. Consistent with our earlier findings, relative to CI expression, the liver mitotype was enriched for multiple FAD-dependent enzymes (**Figure S2G)** and was also enriched for CII by 2.1-fold relative to brain (Figure 2K), confirming pathway-level patterns at the level of single genes.

#### Multivariate mitotype specialization

To systematically describe and quantify multivariate patterns of mitochondrial specialization across multiple functional domains, we took two different approaches. First, we compared all MitoPathways between brain and liver tissues to determine which pathways exhibit higher expression in one tissue relative to the other, and by what magnitude, using ranked log_2_-fold-change analysis (**Figure S4A**). We then visualized two most strongly enriched MitoPathway pairs higher in liver (relative to brain) or vice versa, using bivariate plots of MitoPathway scores. Compared to all other tissues, the liver mitotype was enriched by 4- to 137-fold (48-fold on average) for *Urea cycle* (the liver is responsible for clearing the body’s ammonia by transforming it to urea^28^). The brain was among other tissues enriched for *Glycerol-3-phosphate shuttle* (G3P, brings reducing equivalents from the cytoplasm to the mitochondrial electron transport chain^33^), a shuttle that appears to be of low relevance to liver mitochondria (**Figure S4B**). The second pair of most differentially expressed MitoPathways were *Tetrahydrobiopterin (BH4) synthesis* (2-30x higher in brain, used as co-factor for neurotransmitter synthesis^34^) and *Serine metabolism* (20-80x higher in liver, used in anaplerosis and lipogenesis^35^) (**Figure S4C**). These two-tissue systematic contrasts highlight mitochondrial molecular specialization related to known tissue physiology.

Our second approach consisted in examining the top enriched MitoPathways across the major axes of mitochondrial specialization (the four poles along the multi-tissue mitotype axes, see Figure 2C). **Figure S4D** illustrates families of MitoPathways coordinately enriched among the i) small intestine, ii) skeletal muscle, iii) liver, and iv) cerebral cortex. Each organ exhibited a coherent pattern of overexpression for a family of biologically-related metabolic pathways. In particular, the liver mitotype was expectedly specialized for multiple complementary anabolic pathways (serine and urea metabolism, heme synthesis), as well as xenobiotic metabolism, in line with the liver’s role in systemic detoxification. In contrast, the skeletal muscle mitotype was specialized in electron transport via Coenzyme Q, the intestine mitotype in vitamin A and proline metabolism, and the brain in CI-related biology (**Figure S4E**), possibly underlying its selective vulnerability to CI-related disease^36^. These largely confirmatory findings exemplify a simple approach to visualize and quantify mitochondrial specialization.

Using this framework, we then specifically examined whether mitochondrial specialization occurs within a critical subsystem of mitochondrial biology: OxPhos. The OxPhos system is generally considered the core of mitochondrial biology as it produces and converts the electrochemical gradient into usable forms of energy such as ATP and NADPH^2,19^. As tissue mitotypes exhibit higher FAO expression, they also express more of OxPhos CI, resulting in a roughly linear relationship between both MitoPathways. However, the brain stood out as expressing unusually high levels of CI relative to FAO (**Figure 3A***, left*). In fact, even relative to OxPhos CII and Complex III (CIII), all brain regions express higher levels of CI (Figure 3A, *middle*). Interestingly, all tissues exhibited the same relative abundance of CI and CIV (Figure 3A, *right*). Mitotype ratios demonstrated that the brain expresses on average 60% more CI relative to CIII (*p*<0.001, Hedges’ g = −5.6). The CI:CIII ratio is 2.5 in the brain vs 1.5 on average across all other tissues (**Figure 3B**). In contrast, the average CI:CIV ratio in the brain is similar to all other tissues (CI:CIV = 0.9, **Figure S5B**). Both the brain and heart overexpress CI relative to all other MitoPathways, highlighting a particularly central role for CI and energy transformation in these tissues. This pattern may also reflect CI’s Na+/H+ antiporter function, which is especially relevant in neurons^37^ (**Figures S5D-E**).

**Figure 3:**
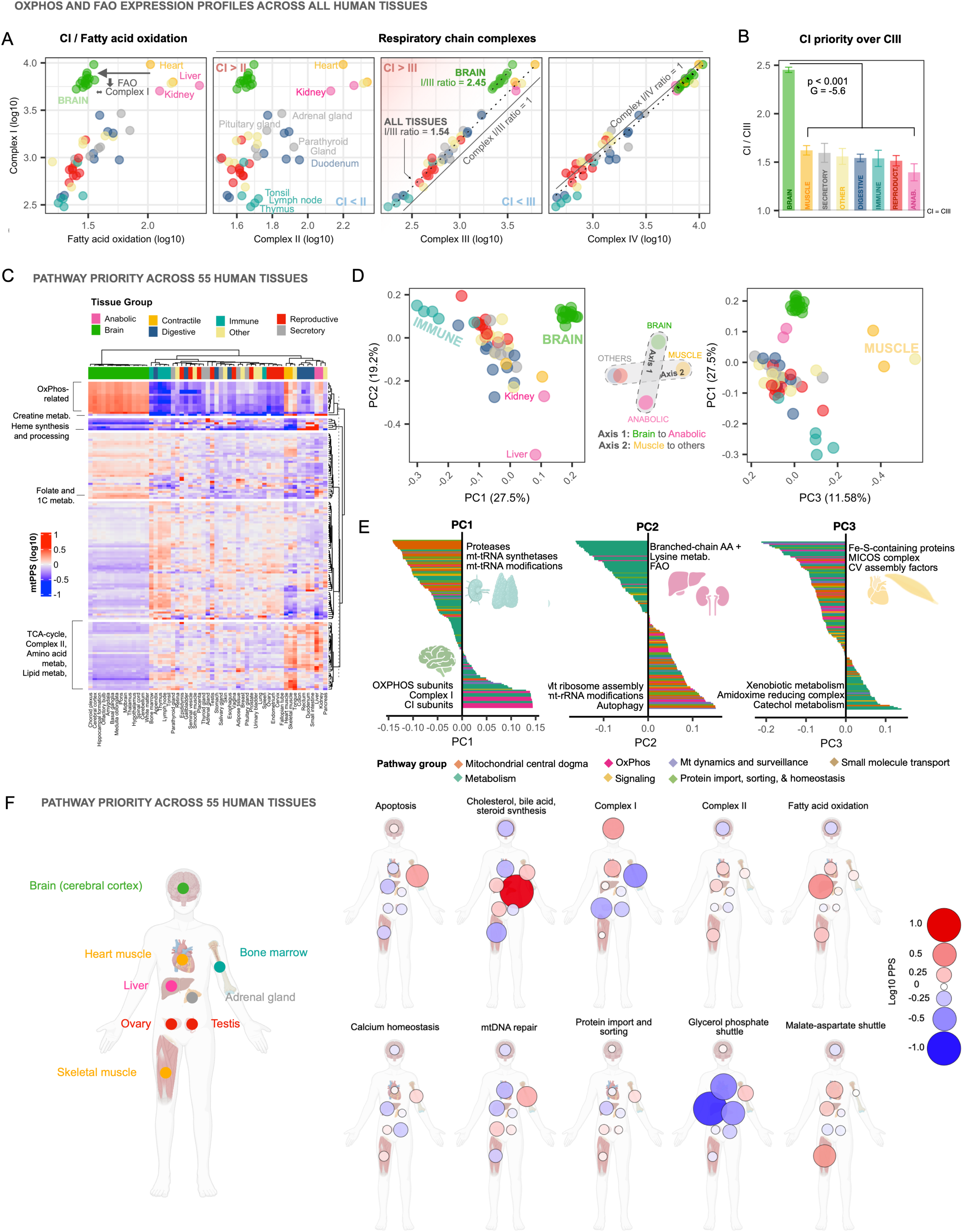
Tissue mitochondrial Pathway Priority Scores. **A)** Bivariate plots of OxPhos and FAO expression profiles in the HPA dataset, illustrating the molecular specialization of mitochondria consistent with known physiological and functional characteristics of each tissue. The solid lines represent identity lines (x = y), the dotted lines indicate linear regression estimates for brain and all other tissues separately. **B)** Mean ratios (±SEM) of CI/CIII pathway scores across tissue groups suggesting that ratio-based approaches can be used to capture mitochondrial pathway profiles, inspiring the development of the mitoPPS framework. Welch’s t-test and hedges’ g for effect size. **C)** Heatmap and hierarchical clustering of mitoPPS in log10 scale in the HPA dataset showing distinct prioritization signatures for different tissues, where a score of 0 represents average priority. Rows are pathways and columns are tissues. For instance, compared to average prioritization across all tissues, the brain prioritizes OxPhos-related pathways (except Complex II) by an extra ∼110% (i.e., MitoPPS value = 2.1), while immune tissues deprioritize OxPhos by - 65% (MitoPPS value = 0.35). Rows are k-means clustered, and column clusters are calculated using the Euclidean distance and the WardD.2 method. **D)** Principal component analysis of mitoPPS shows that PC1 separates immune and brain tissues, while PC2 separates anabolic tissues, and PC3 separates contractile muscle tissues. **E)** Loadings for PC1, 2, and 3. The top 3 pathways are highlighted, and the pathway loadings are color-coded by level1 MitoPathway annotations. Tissue symbols indicate the tissue group mostly associated with each PC. **F)** mitoPPS across the human body for a subset of 8 tissues, and 10 MitoPathways. mitoPPS on log10 scale, with 0 being average priority (white, small circle), 1 being higher priority (red, large circle), and −1 indicating low priority (blue, large circle). Abbreviations: PC, principal component; AA, amino acid

To summarize, so far we have shown that two-tissue contrasts, multi-tissue and multi-pathway signatures, as well as MitoPathway ratios, confirm and quantify the magnitude of mitochondrial specialization across physiologically divergent human organs and tissues.

#### Mitochondrial pathway prioritization scores (mitoPPS)

While informative, the approach based on simple MitoPathway scores described above is limited in three ways: First, it is influenced by overall mitochondrial gene expression levels. As a result, tissues with inherently high mitochondrial content (e.g., kidney, heart, liver, muscle, brain) consistently produce the highest scores, with values spanning several orders of magnitude compared to low-mitochondria tissues. Second, the scores lack directionality and do not sufficiently quantify “deprioritization” of pathways, such as the G3P shuttle in the liver (see Figure S2B). Third, MitoPathway scores are only in relative terms across samples within a dataset (i.e. higher in tissue A relative to tissue B or the dataset average). They cannot be expressed on an absolute scale, which prevents comparisons across datasets and modalities, and pathways.

To resolve these limitations, we developed an approach that integrates MitoPathway *ratios* rather than absolute scores between all 149 pathways across 55 tissues. This produced ∼1.2 million comparisons, with values spanning eight orders of magnitude. To make the resulting metric independent of absolute expression levels and tissue mitochondrial content, we generated corrected ratio-based scores that quantify the extent to which each MitoPathway is prioritized in a given sample, relative to all other MitoPathways and samples. The resulting *mitochondrial Pathway Prioritization Score (mitoPPS)* and underlying computational structure is illustrated in **Figure S6** with CI in brain and liver as example.

Briefly, the mitoPPS approach consists in normalizing each MitoPathway expression to each of the other 148 pathways. The resulting normalized ratios are then averaged for each pathway and tissue. This yields *the fraction of the mitochondrial transcriptome devoted to each MitoPathway relative to other pathways*. It thus reflects how much a given function or behavior is “prioritized”, or how much resources a given cell/tissue tries to invest in a given MitoPathway (**Figure S6A-B**). The resulting simple linear scalar is an interpretable metric – the higher the score, the more prioritized is a given MitoPathway. The mitoPPS scores are also comparable across samples and datasets, provided they offer similar coverage of the underlying genes and proteins. Importantly, the mitoPPS scores generated do not suffer from the overall mitochondrial abundance bias. The mitoPPS and the z-scored MitoPathway expression scores were significantly correlated, particularly when examining non-mtDNA (Spearman’s ρ_non-mtDNA_=0.31, *p*<0.001) vs mtDNA-encoded MitoPathways (ρ_mtDNA_=0.73*, p*<0.001) separately (**Figure S7**).

#### Mapping mitotype specialization across tissues using mitoPPS

Applied to the multi-tissue dataset, the mitoPPS produces an interpretable metric of MitoPathway prioritization for all 55 tissues (**Figure 3C**). Compared to the raw MitoPathways scores, the dynamic range for mitoPPS scores spans three orders of magnitude (∼1,000-fold, ∼0.01-14), likely reflecting a more realistic physiological range of mitochondrial specialization across the human body than the four-order-of-magnitude range observed for raw MitoPathway scores.

Some striking findings include: i) Brain tissues were most enriched for *OxPhos* whereas immune cells had the lowest *OxPhos* expression (6-fold difference, *p*<0.0001). ii) *Heme synthesis and processing* dominated in the liver, as well as some, albeit not all, digestive tissues (16-fold difference, *p*<0.0001). iii) *Creatine metabolism* was most prioritized in both anabolic and contractile muscle tissues, as well as pancreas (10-fold difference, *p*<0.001). and iv) *Folate and 1C metabolism* was most prioritized among brain tissues (except the choroid plexus) and liver (2.3-fold difference, *p*<0.0001). The quantitative distribution of mitoPPS scores underscores the interdependence of organ systems^38^, arising from the apparently exclusive specialization of mitochondria for some functions in specific tissues (discussed further below).

Using mitoPPS as input, the multi-tissue data were projected into PCA space, where the first three components explained ∼58% of the variance (**Figure 3D**), broadly matching the results from gene-based PCA (Figure 2E). The brain mitotype again emerged as strikingly distinct, opposed by anabolic tissues (the liver in particular), muscles, and other tissues. The conservation of the two major axes observed in the gene-based PCA suggests that mitoPPS does not introduce artificial bias, while adding quantitative resolution and greater interpretability. From the PCA loadings, we identified the top pathway categories driving each axis (**Figure 3E, Supplemental File 2)**. *PC1* reflects a polarization from OxPhos (positive, *magenta*) to mitochondrial central dogma-related pathways (negative, *orange*). *PC2* contrasts metabolic pathways (negative, *teal*) from a mixed set of pathways (positive). And *PC3* contrasts metabolism-related pathways (positive, *teal*) from a mixed set of pathways including OxPhos and mitochondrial central dogma (negative, *magenta* and *orange*, respectively).

This analysis highlights clear tissue-specific priorities. Brain and immune mitotypes diverge sharply on PC1 (OxPhos is high in brain mitochondria, mtDNA-related pathways are enriched in immune mitochondria). Yet, both deprioritize metabolism compared to liver, which emphasizes lipid and amino acid metabolism. Muscle mitochondria, by contrast, were most distinct along PC3, driven by their prioritization of iron-sulfur proteins, the MICOS complex, and CV assembly factors.

**Figures 3F** and **S8** summarize the prioritization of 10 major mitochondrial pathways across eight vital organs. The adrenal gland mitotype strongly prioritizes Cholesterol, bile acid, and steroid synthesis pathways (14.6-fold over other tissues) consistent with its specialized role in steroidogenesis (adrenal mitochondria being the major site of glucocorticoid hormone synthesis^39^). Co-prioritization of multiple mitoPPS across organ systems, highlighted in the vertical dendrogram of Figure 3C (more details in **Figure S7A**) further shows how adrenal mitochondria specialize in complementary endocrine pathways, such as Vitamin D metabolism (19.4-fold), and, to a lesser extent, in other pathways required for redox-based biosynthetic reactions (e.g. Fe-S biosynthesis, 1.85-fold). The liver similarly displays a distinct mitochondrial signature and again shows low reliance on the G3P shuttle, reinforcing the pattern observed in the brain-liver comparison (Figure S4A-B). This extends to other oxidative tissues: heart and skeletal muscle favor the malate-aspartate shuttle over G3P, reflecting their shared reliance on NADH-linked oxidation and redox-efficient transport through CI. Bone marrow, in contrast, favors the G3P shuttle over MAS despite its glycolytic profile characteristic of hematopoietic stem and progenitor cells^40,41^. This likely reflects the need for redox flexibility and partial electron transport independent of CI, allowing hematopoietic and progenitor cells to sustain cytosolic NAD⁺ regeneration and electron flux during proliferation and differentiation^42,43^.

In summary, mitoPPS captures a coordinated mosaic of mitochondrial specializations across human tissues, reflecting both the division of metabolic labor and the inter-tissue coupling required to sustain organismal homeostasis.

#### In vitro plasticity of mitochondrial pathway prioritization

Many of the mitotype differences described above are likely driven by robust differences in cell-type specific metabolic, biosynthetic and signaling requirements inherent to organismal physiology ^1^. To determine the extent to which mitochondrial phenotypes can exhibit plasticity *in a given cell type*, we used a controlled *in vitro* monoculture system of primary human dermal fibroblast. The *Cellular Lifespan Study* dataset includes RNAseq data from multiple donors (n=8) exposed to pharmacological, metabolic, and endocrine challenges (n=13 treatment conditions), longitudinally monitored for up to 9 months (40 passages)^44^. Compared to the GTEx dataset that examines mitotypes from different organs in their normal physiological states, this system allows us to ask in a more definitive manner which MitoPathways are obligatorily (i.e., genetically) co-regulated, and which subsets of mitochondrial functions can become uncoupled in response to stress or during progression toward cellular senescence.

#### Longitudinal mitotype trajectories

To interrogate how mitotype profiling captures dynamic, time-resolved mitochondrial recalibrations, we first applied our mitoPPS analysis to longitudinal transcriptomic data from untreated primary fibroblasts derived from the three most deeply phenotyped donors (Healthy Controls: HC1-3). We previously confirmed in this model that time in culture, or the number of passages, is associated with canonical hallmarks of cellular senescence including progressive telomere shortening, increased epigenetic clocks, and heightened expression of senescence-associated markers^45–47^. As shown in **Figures 4A-B**, RNA levels of senescence indices (CDKN2A, CDKN1A, TP53) rise while proliferation markers (KI67, TOP2A, RRM2) decline, confirming the activation of senescence-associated programs. This provides an ideal setting for assessing mitochondrial recalibrations associated with the transition into senescence.

**Figure 4:**
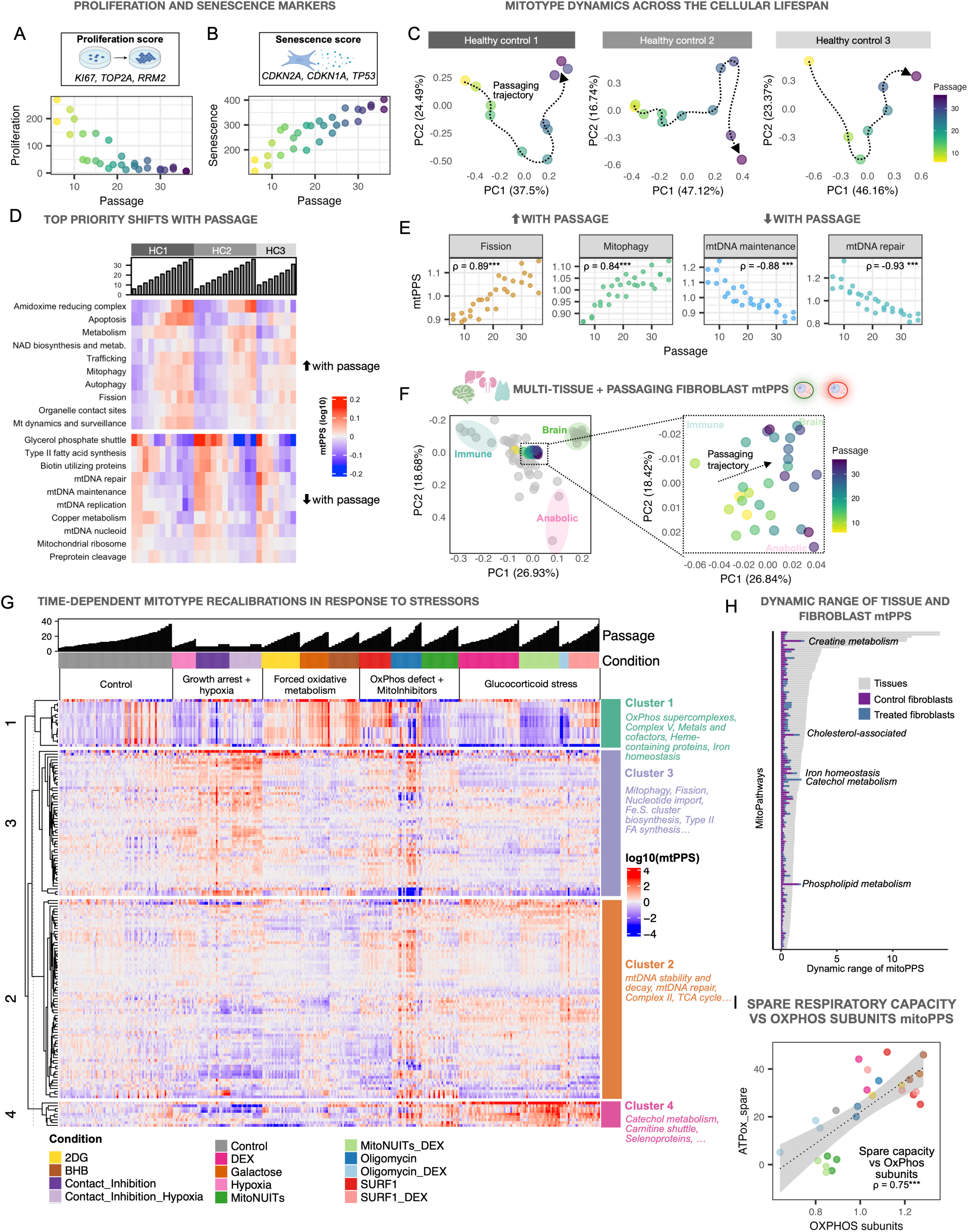
Mitochondria dynamically recalibrate in response to experimental perturbations. **A)** The proliferation index (average expression of genes *KI67, TOP2A, and RRM2*) is downregulated and **B)** the senescence index (*CDKN2A, CDKN1A, and TP53*) is upregulated in healthy control fibroblasts cultured over 30+ passages. **C)** Principal component analysis of MitoPathway mitoPPS from three healthy controls. Dotted lines indiciate individual aging trajectories, suggesting a gradual MitoPathway priority shift with cellular age. **D)** Heatmap of top ten up – and downregulated mitoPPS (log10) with age in three healthy controls. Columns represent samples sorted by cell line and ascending passage. **E)** Scatter plot and spearman’s ρ of two pathways that become prioritized (Fission, Mitophagy, *left panel*) and deprioritized (mtDNA maintenance, mtDNA repair, *right panel*) with age. **F)** PCA of the multi-tissue Human Protein Atlas mitoPPS with the immune, brain, and anabolic axes, and Cellular Lifespan Study mitoPPS (left panel). Each dot represents one tissue (grey) or fibroblast sample from healthy controls (color gradient by passage). The right panel shows the same PCA as in the left panel, highlighting the fibroblast samples. The pathways loading most strongly on each PC for each donor are listed in Supplemental File 2. **G)** Heatmap of mitochondrial recalibrations in response to experimental perturbation. Rows represent MitoPathways and columns represent samples, sorted by experimental condition and ascending passages. **H)** Dynamic range of tissue and fibroblast mitoPPS. For tissues, the dynamic range was calculated across 55 human tissues using the delta between the lowest and highest mitoPPS. For control fibroblasts, the dynamic range was determined along the lifespan trajectories show in Figure 6C from 3 cell lines. The treatment fibroblast dataset was first averaged across cell lines and conditions to remove the aging effect, and the dynamic range was determined using the delta mitoPPS from lowest to highest. **I)** Bivariate plot and spearman correlation coefficient of spare respiratory capacity and OxPhos subunits mitoPPS across all experimental conditions.

The mitoPPS-based mitotype signatures across all timepoints projected in PCA space revealed that mitochondrial remodeling towards senescence proceeds in at least two phases (**Figure 4C**). For example, the mitotype of HC1 initially shifts downward (along PC2) and rightward (along PC1) over the first 21 passages (∼100 days), then gradually ascends along PC2 while remaining relatively constant on PC1 at later passages. Interestingly, each donor exhibited a relatively unique trajectory, possibly mirroring inter-individual heterogeneity ^48^. We then asked which mitoPPS changed most consistently with increasing cellular senescence. The top up- and downregulated pathways (**Figure 4D)** revealed coordinated, time-dependent remodeling. Among the most upregulated pathways over time were mitochondrial fission and mitophagy involved in mitochondrial quality control (**Figure 4E**). In contrast, among the most downregulated pathways were mtDNA-related pathways, such as maintenance and repair, the same pathways associated with immune mitotypes (see PCA loadings in Figure 5E).

**Figure 5.**
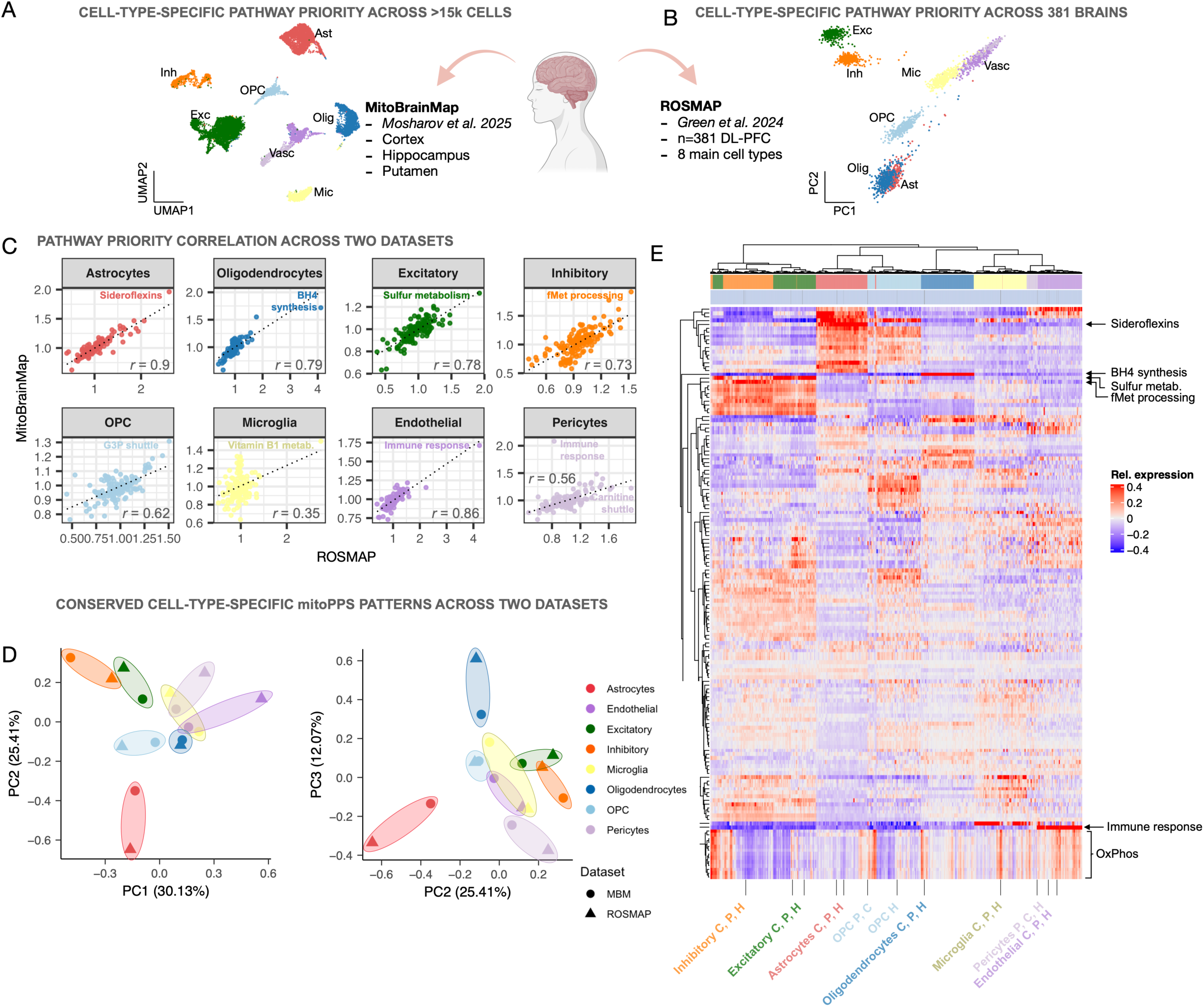
Mitotyping reveals reproducible cell-type-specific mitochondrial specialization across independent human brain datasets. **(A)** UMAP representation of grey matter cells from the MitoBrainMap single nuclei RNA sequencing dataset^49^ used to compute mitochondrial pathway priority scores (mitoPPS) across major neuronal, glial, and vascular populations. **(B)** Principal component analysis (PCA) visualization of the ROSMAP cohort (Green et al., 2024; *n* = 381 dorsolateral prefrontal cortex samples) used for mitoPPS **(C)** Cross-dataset correlation of MitoPathway prioritization between MitoBrainMap and ROSMAP demonstrates high reproducibility (average Pearson’s *r* = 0.7), with the strongest correspondence observed in astrocytes (*r* = 0.9). Top-prioritized pathways included sideroflexins in astrocytes, tetrahydrobiopterin (BH₄) synthesis in oligodendrocytes, sulfur metabolism in excitatory neurons, and *fMet* processing in inhibitory neurons. **(D)** PCA of MitoPathway scores shows clustering by cell type rather than dataset along principal components (PC) 1, 2, and 3, indicating convergence of mitotype profiles across independent cohorts. **(E)** Hierarchical clustering of pathway-level scores reveals conserved, cell-type–specific mitochondrial signatures, including consistent top-prioritized pathways across datasets. Abbreviations: Exc, excitatory neurons; Inh, inhibitory neurons; OPC, oligodendrocyte precursor cells; Vasc, vascular cells; Ast, astrocytes; Olig, oligodendrocytes; Mic, microglia; BH4, tetrahydrobiopterin; G3P, glycerol-3-phosphate; MBM, MitoBrainMap; C, cortex; H, hippocampus; P, putamen; DL-PFC, dorsolateral prefrontal cortex;

This finding prompted us to test whether mitoPPS from the longitudinal fibroblast dataset could be integrated with the multi-tissue dataset. This integration is possible given that both mitoPPS datasets are unitless and internally normalized to other MitoPathways, representing a major strength of this approach.

The fibroblast mitoPPS clustered near the center of the tissue distribution (**Figure 4F**). Projecting fibroblasts into the multi-tissue space revealed that with cell senescence, mitochondria shifted along PC1 away from immune tissues (which upregulate mtDNA-related pathways) toward the more catabolic brain mitotype. Consistent with this, OxPhos-related mitoPPS (CI, CIII-CV, upregulated in brain mitotypes) increased with passage in healthy, untreated cells (**Figure 4G** left side of the heatmap). At a high level, this pattern across the cellular progression to senescence may indicate a shift in priority from a proliferative/signaling mitotype (immune tissues) towards a less versatile, energy-focused senescent mitotype. These findings showcase mitoPPS-based mitotyping from bulk RNAseq in cultured cells, and its easy integrability with a multi-tissue human reference.

#### Mitotype recalibrations to targeted metabolic perturbations

We next examined how much plasticity the human fibroblast mitotype can exhibit in response to various metabolic perturbations: OxPhos inhibition with low dose oligomycin, genetic mutations in the SURF1 gene that impairs basal OxPhos and upregulates glycolysis^47^, hypoxia (3% O_2_ compared to ambient 21%), and long-term growth arrest (up to 140 days in contact inhibition without passaging). Metabolic manipulations that influence mitochondrial carbon substrates availability included: i) galactose without glucose or ii) 2-deoxyglucose (2-DG) that directly inhibits glucose metabolization (glycolysis, pentose phosphate pathway); iii) 2-beta-hydroxybutyrate (BHB), a ketone body oxidized in mitochondria; and iv) a combination of three mitochondrial nutrient uptake inhibitors (NUITs: UK5099, Etomoxir, BPTES) that block mitochondrial pyruvate, medium- and long-chain fatty acid, and glutamine transport, starving the Krebs cycle of carbon substrates (see^44^ for details). Together, these chronic experimental conditions applied for up to 9 months constitute a robust test whether and how much the mitotype of a given cell type can recalibrate.

For each timepoint and cell line, we calculated mitoPPS relative to controls across all treatments (**Figure 4G** and **S9**). Multi-treatment data was integrated into the multi-tissue mitoPPS reference (**Figure S9D-E**). Overall, treatments induced coordinated effects across multiple MitoPathways. Glucose availability reduction (2-DG, Galactose) or BHB forced the expected shift toward OxPhos-related pathways (Cluster 1 in Figure 4G) and occupied a similar PCA space shifted towards the oxidative and catabolic brain-like mitotype. In comparison, Galactose-treated fibroblasts prioritized CI by +18%, while the cerebral cortex prioritized CI by +163% relative to all other tissues (**Figure S10A**). On the other hand, a combined treatment of the synthetic glucocorticoid Dexamethasone and NUITs induced a shift toward an anabolic mitotype along PC2, with modest FAO increases (+11-13%), far less than the liver (+216% **Figure S10B**). These results confirm expected treatment-specific patterns of mitochondrial plasticity, but at magnitudes substantially smaller than tissue-level differences.

Fibroblast mitotype plasticity thus spans a narrower dynamic range than observed across tissues (**Figure 4H**). Nonetheless, some perturbations produced pathway-specific changes approaching or exceeding inter-tissue variation (**Figure S11B-C**). Even within a single cell type, mitochondria can therefore undergo marked and highly plastic remodeling, as captured by mitoPPS.

#### Linking MitoPPS to bioenergetic function

Because transcriptomic signatures do not necessarily reflect functional activity, we next asked what mitoPPS captures in bioenergetic terms and how pathway prioritization relates to measurable mitochondrial performance. To do so, we correlated mitoPPS values across all treatment conditions with Seahorse-derived bioenergetic readouts, including OxPhos- and glycolysis (i.e. lactic-fermentation)-linked ATP rates at baseline (ATPox and ATPglyc) and under maximal respiration (ATPox_max, ATPox_spare, measured following FCCP uncoupling), as well as proton leak, coupling efficiency, and non-mitochondrial respiration (i.e. oxygen consumption after complete ETC inhibition). This analysis links mitoPPS to real-time cellular bioenergetic output, allowing us to interpret mitoPPS scores in terms of bioenergetic mitochondrial behavior.

At baseline, ATPox showed no overall correlation with OxPhos-related mitoPPS, indicating that prioritizing OxPhos pathways does not necessarily translate into higher instantaneous ATP generation. Instead, ATPox was significantly negatively correlated with mitochondrial central dogma pathways (e.g. replication, transcription, translation, ρ = −0.76, p_adj_<0.001, **Figure S11D** and **S11F**), but positively correlated with the cholesterol-associated pathway (ρ = 0.77, p_adj_<0.001, **Figure S11E**-F). In contrast, spare respiratory capacity (i.e. the ability to increase ATP production when challenged) showed the strongest positive correlation with OxPhos mitoPPS across all treatment groups (**Figure 4H**). This contrast suggests that mitoPPS does not primarily reflect steady-state ATP production but instead reflects the capacity to mobilize OxPhos when required. In other words, in this system mitoPPS appears to capture *bioenergetic preparedness* rather than ongoing flux.

#### Cell-type resolution mitotyping in the human brain across datasets

To evaluate whether mitotyping can be reproducibly applied in different cell types, we next applied mitoPPS analysis to two independent pseudobulk single-nucleus RNA-seq datasets of the human cortex: data from our previously published MitoBrainMap atlas^49^ (**Figure 5A**) and the ROSMAP cohort^50^ (**Figure 5B**). Despite differences in cohorts, brain regions, and technical processing, the mitotypes for major neuronal, glial and vascular classes were highly correlated across the two datasets (**Figure 5C**; average Pearson’s r = 0.7). The strongest correlation was observed for astrocytes (r = 0.9) whereas neuronal subtypes showed greater variability (r = 0.73 – 0.79), likely reflecting regional heterogeneity and donor-specific influences.

The top-prioritized mitochondrial pathways were consistently recovered in both resources, underscoring conserved, cell-type specific specializations that transcend dataset-specific artifacts. Astrocytes robustly prioritized sideroflexin pathways, oligodendrocytes were enriched for tetrahydrobiopterin (BH4) synthesis, excitatory neurons emphasized sulfur metabolism, and inhibitory neurons prioritized fMet processing. Pericytes were the only notable exception, likely reflecting their low abundance in both datasets. Despite this, the same top pathways were recovered in both datasets, although their prioritization magnitudes differed (Figure 5C). For instance, the BH4 synthesis score in oligodendrocytes was roughly 2-fold higher in MitoBrainMap but about 4-fold higher in ROSMAP brains compared to average. After correcting for these scale differences, all cell types converged in PCA space (PC1-3, **Figure 5D, S12B-C**) and clustered by cell type rather than dataset (**Figure 5E** and **S12A**), supporting the conservation of mitochondrial pathway priority score patterns across independent cohorts.

Residual differences in magnitude likely reflect biological rather than technical variation, including differences in cell subpopulation composition, disease state, and age^50^ (ROSMAP encompasses an aged population, whereas the MitoBrainMap donor was a 54-year-old apparently healthy individual), plus broader cortical coverage in ROSMAP relative to MitoBrainMap. Consistent with this, we also observed substantial inter-individual variability within the ROSMAP cohort itself, and top-prioritized pathways remained prominent even at the population level (Figure 5E), underscoring that mitotyping captures both conserved and context-dependent mitochondrial features.

Together, these results demonstrate that mitotyping yields robust, biologically coherent, and reproducible mitochondrial signatures across pseudobulked single-nucleus RNAseq datasets, yielding stable estimates of cell-type-specific mitochondrial specializations in the human brain. At the same time, the observed inter-individual variability in pathway prioritization in the ROSMAP cohort suggests that cell-type specific mitochondrial programs are not rigid but can recalibrate dynamically *in vivo*.

### Single-cell mitotyping along spermatogenesis

#### Capturing mitochondrial reprogramming along a developmental trajectory

To further challenge our method and explore its sensitivity to known developmental metabolic needs, we next asked whether mitoPPS can resolve continuous trajectories of mitochondrial specialization in human spermatogenesis as a model system. We used this model as it follows a well-defined sequence of germ cell states, from spermatogonial stem cells (SSC) and differentiating spermatogonia, to spermatocytes and spermatids^51^ (**Figure 6A**). This process is accompanied by coordinated shifts in mitochondrial morphology, metabolism, and bioenergetic output^51–53^, ultimately producing mature sperm with densely packed and highly efficient mitochondria that provide the energy required for motility and fertilization^54^. We therefore reasoned that this developmental continuum provided an ideal testbed for mitotyping along a continuous cell lineage, allowing us to test whether mitoPPS scores capture the progressive reorganization of mitochondrial programs among the transcriptional “noise” linked to other cellular and developmental changes.

**Figure 6.**
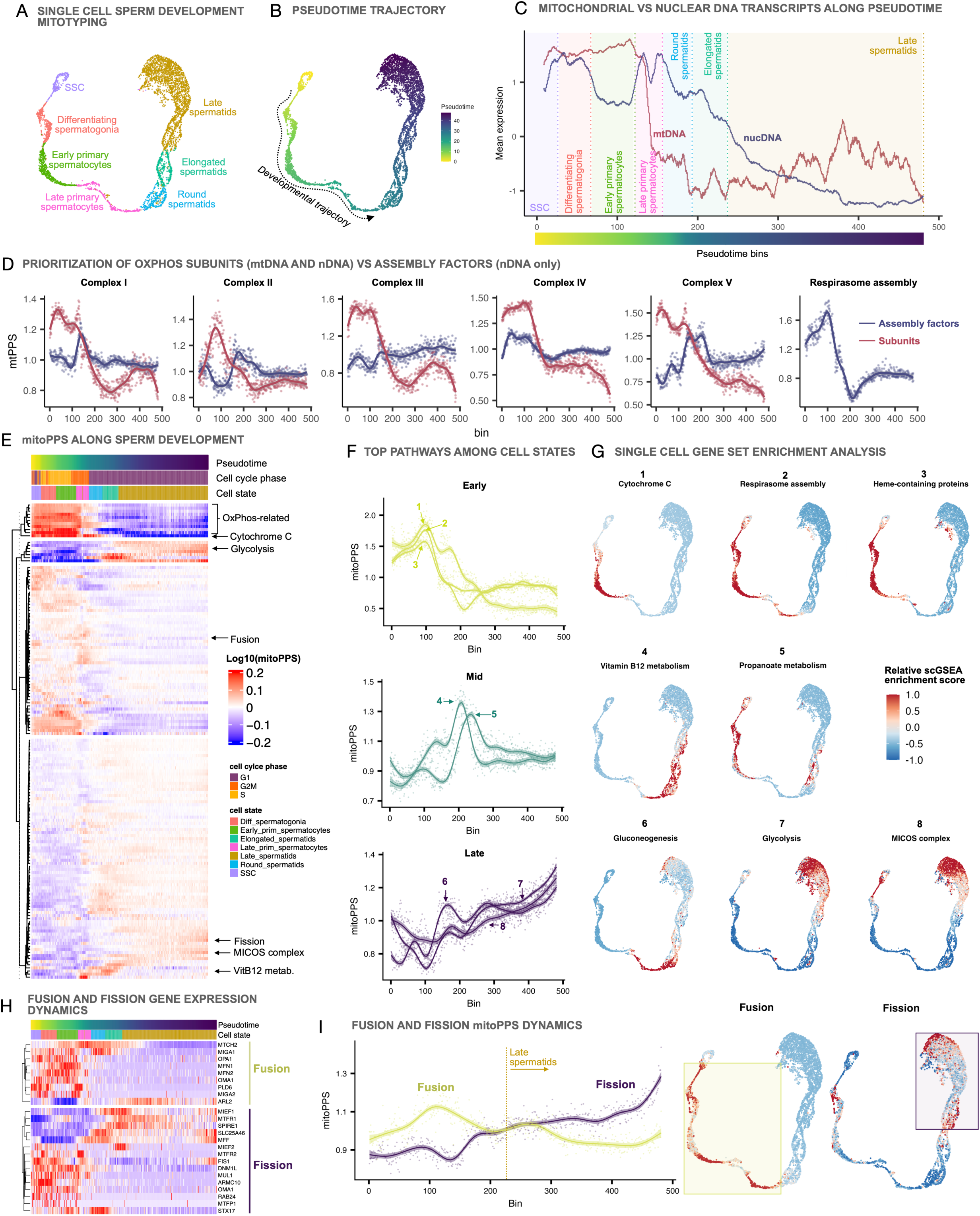
Single-cell mitotyping reveals stepwise mitochondrial specialization during human spermatogenesis. **(A)** Schematic of spermatogenic differentiation, from spermatogonial stem cells (SSCs) to mature spermatids in a human testis single-cell RNA-seq dataset ^55^highlighting major developmental stages. **(B)** Pseudotime trajectory reconstructed the data shown in A) illustrating the continuous progression of germ cell states. **(C)** Dynamics of mitochondrial DNA (mtDNA) and nuclear DNA (nDNA)-encoded transcripts along pseudotime. The solid lines indicate smoothed mean expression profiles along pseudotime, using the scaled rolling mean (window size = 80 cells) per genome. Vertical dotted lines mark cell stages. mtDNA transcripts decline sharply following the last mitotic division, followed by a gradual reduction in nDNA-encoded mitochondrial transcripts. **(D)** mitoPPS analysis of oxidative phosphorylation (OxPhos) complexes I–V and respirasome assembly reveals inverse regulation of subunit and assembly factor expression, especially in CII and CV. **(E)** Heatmap of MitoPathway prioritization along pseudotime shows temporal shifts in mitochondrial pathway prioritiy, including early OxPhos enrichment and late-stage metabolic rewiring. **(F)** Top prioritized pathways across early, mid, and late stages. Early enrichment of OxPhos-related and cytochrome *c* pathways; mid-stage prioritization of vitamin B12 and propanoate metabolism; and late activation of gluconeogenesis, glycolysis, and the MICOS complex, indicating cristae remodeling and mitochondrial maturation. **(G)** Single-cell gene set enrichment analysis (scGSEA) validates stage-specific pathway activity (FDR < 0.05) and visualizes mitotype transitions across spermatogenesis on the single cell level. Scores are relative across pseudotime. **(H–I)** Expression and mitoPPS dynamics of mitochondrial fusion and fission pathways reveal early fusion prioritization, followed by progressive fission activation during later stages, consistent with the transition to compact, specialized mitochondria in mature sperm. Abbreviations: SSC, spermatogonial stem cell;

To create a continuous time axis, we computed a pseudotime trajectory from a publicly available testis single cell RNA seq dataset (^55^, **Figure 6B**) from the human protein atlas^22^. Because mitochondrial DNA (mtDNA) is destined to degrade in sperm development^56–60^, we first examined mtDNA vs nuclear DNA (nDNA) gene expression dynamics. We calculated average expression scores for all mitochondrial genes encoded by either genome (n = 13 mtDNA, n = 1094 nDNA genes). As expected, mtDNA-encoded transcripts declined sharply following the last mitotic division (early to late primary spermatocytes). This was followed by a subsequent decrease in nDNA encoded mitochondrial transcripts (**Figure 6C** and **S14**). Since mtDNA encodes exclusively core respiratory chain subunits while nDNA encodes the assembly factors, we next examined their dynamics using mitoPPS. Given the sparsity and noise typical of single-cell data, we binned cells along pseudotime (10 cells per bin) and computed mitoPPS for each bin (see Figure 6C for stage-bin-pseudotime correspondence). Interestingly, assembly factor expression increased as subunit expression decreased, most prominently in Complex V (**Figure 6D**). Notably, respirasome assembly (nDNA-encoded assembly of supercomplexes) is highly prioritized in early primary spermatocytes (mitoPPS > 1.6) and declines sharply thereafter, coinciding with subunit downregulation, potentially reflecting supercomplex assembly events in sperm, a largely understudied phenomenon.

#### Stage-specific metabolic programs and mitochondrial remodeling

We then extended our analysis to all MitoCarta pathways and included glycolysis, which plays a key role in sperm energy metabolism ^54,61^, as well as its anabolic counterpart, the pentose-phosphate pathway. Mitochondrial prioritization markedly changed after differentiation was complete (**Figure 6E**). Early stages were dominated by strong OxPhos and associated signatures, including cytochrome c, consistent with its unique role during spermatogenesis^62^ (**Figure** 6D, **6F** *top panel***, and S15A**). In mid stages, vitamin B12 and propanoate metabolism pathways were prioritized, potentially indicating a transient shift toward co-factor dependent anaplerosis that supports TCA cycle replenishment (Figure 6F *middle panel*). Vitamin B12 serves as a cofactor for methylmalonyl-CoA mutase, which links propanoate metabolism to succinyl-CoA production and sustained mitochondrial ATP generation. In contrast, mid to late stages prioritized gluconeogenesis, followed by glycolysis, suggesting a terminal phase of metabolic rewiring (Figure 6F *bottom panel***, Figure S15B**). The brief gluconeogenic activation may recycle carbon to sustain glycolytic flux, while glycolysis itself provides localized ATP within the developing flagellum^61^.

To validate these findings, we performed single-cell gene set enrichment analysis (scGSEA^63^), which confirmed stage-specific pathway activity (false discovery rate <0.05). Averaging results across multiple scGSEA runs and random seeds ensured robustness, and the results recapitulated mitotype transitions along spermatogenesis at single cell resolution (**Figure 6G**). Importantly, mitoPPS and scGSEA are methodologically independent yet yielded consistent patterns. Yet, unlike enrichment-based methods, in which pathway scores cannot be directly compared between each other, mitoPPS produces a quantitative, cross-comparable metric that also captures relative effect sizes.

#### mitoPPS links mitochondrial remodeling to metabolic state transitions during spermatogenesis

Mitochondrial architecture and metabolic remodeling are closely coupled throughout spermatogenesis. In SSCs, mitochondria are small and spherical with orthodox cristae, reflecting a glycolytic and low-OxPhos metabolic profile typical of undifferentiated cells. As SSCs differentiate into spermatogonia and progress toward early primary spermatocytes, mitochondria elongate and cluster, coinciding with elevated OxPhos activity to meet the energetic demands of meiotic entry. In round spermatids, mitochondria fragment, together with a metabolic shift back toward glycolysis, while in elongated and late spermatids, fission predominates to form the compact mitochondrial sheath characteristic of mature sperm ^52,64,65^.

These morphological and metabolic transitions were reflected in our pathway priority scores. The progression from SSCs to differentiating spermatogonia was accompanied by a decline in glycolytic prioritization and a rise in OxPhos and fusion. When glycolysis reached its lowest point, fusion priority peaked in early primary spermatocytes, aligning with maximal OxPhos priority (**Figures S15C-D**). During spermatid stages, glycolysis and fission increased together, consistent with the transition from a dynamic mitochondrial network to the compact, organized architecture of mature sperm. Consistent with structural maturation, the MICOS complex, which shapes mitochondrial cristae, was highly prioritized (**Figure S15E**), likely marking final inner-membrane reorganization and assembly of efficient mitochondria in spermatids.

Collectively, these results reveal a stepwise program of mitochondrial specialization along spermatogenesis and demonstrate that mitoPPS effectively captures and interprets these transitions. Thus, this method provides a powerful tool to investigate how mitochondrial identities are specified, remodeled and stabilized across developmental lineages, and to interpret mitochondrial states from single-cell data.

## Discussion

Different cell types share the same genetic hardware but run specialized transcriptional programs that subserve specialized metabolic needs. Altogether, this sustains organismal survival. To capture the remarkable diversity and plasticity of mitochondria underlying these processes, we developed an integrative computational framework that extracts mitochondrial pathway prioritization scores (mitoPPS) from transcriptomic data using all to date described mitochondrial pathways. By applying this mitotyping pipeline to systematically and quantitatively define mitochondrial phenotypes (i.e., mitotypes) across mouse and human tissues, as well as in cultured human fibroblasts, the human brain, and across the developmental trajectory of human sperm, we demonstrate that mitochondria from the same organism – and even the same cell type – can adopt highly distinct molecular profiles that change dynamically over time and in response to metabolic challenges. The mitotyping approach, accessible in the associated code, equips both non-experts and experts with a portable approach to navigate and interpret the multifaceted, dynamic nature of mitochondrial biology.

Mitotyping differs from traditional gene set enrichment analyses in two main ways. First, it includes a detailed curated inventory of interpretable mitochondrial functions (e.g., TCA cycle, CI assembly) rather than broader compartmental labels (e.g., mitochondrial cristae or matrix). Second, since it is not confounded by overall mitochondrial transcript abundance, it can be readily integrated across datasets. Our findings raise the question whether some observed molecular signatures reflect stable mitotypes or dynamic mitochondrial states – similar to transient “activated”, “progenitor”, “differentiated”, “aged”, and other cellular states^66,67^. In tissues as in cells, mitotypes likely reflect a combination of both, especially since different mitotypes can co-exist within sub-compartments of the same cell^7,68,69^. Our approach further enables sensitive tracking of dynamic mitochondrial state changes in a cell-type specific manner and importantly, at single cell resolution. This offers a new and powerful tool to quantify changes on the single cell level.

Some limitations should be noted. First, mitoPPS-based enrichment reflects the activation of cellular programs (i.e., what the cell is attempting to accomplish), rather than the actual functional capacity of its mitochondrial population. Future studies should integrate paired RNAseq, proteomics, fluxomics, imaging, and other modalities to map the temporal relationship between the initial shift in transcriptional mitotypes and the resulting protein-level or functional adaptations. Second, mitotype shifts also could occur independent of transcription, suggesting that mitotyping captures only part of the full mitochondrial regulatory spectrum. Nevertheless, this framework can help identify potential “bottlenecks”, i.e. pathways that receive disproportionate cellular attention or resource allocation due to stress, defects, or compensatory demands. Finally, a minority of MitoPathways contain relatively few genes, which may reduce the robustness of their mitoPPS estimates. Pathway curation remains an ongoing challenge. While our approach leverages MitoCarta, this coverage can be expanded as new mitochondrial processes are characterized. For instance, in the spermatogenesis mitotyping, we incorporated two non-mitochondrial yet metabolically essential pathways (glycolysis, pentose-phosphate pathway), illustrating how the framework can flexibly accommodate emerging knowledge. Overall, the proposed quantitative, fine-grained, scalable approach to profiling mitotypes provides a foundation for systematically quantifying and interpreting the multivariate landscape of mitochondrial biology in cells and whole organisms.

## Author contributions

A.S.M. and M.P. designed and conceptualized the study. A.S.M. generated the figures, and wrote the analysis code. A.S.M., M.P., J.A.E., D.K., J.D., and G.S. interpreted the results. J.D., D.K., G.S., and C.T. assisted with data analysis. P.L.D.J and V.M. contributed data and provided methodological guidance. A.S.M. and M.P., drafted the manuscript. All authors provided feedback on the manuscript.

## Acknowledgements

The authors are grateful to Michio Hirano, Alexander Behnke, Irla Belli, Rose Underberg, and members of the Picard lab for helpful discussions and input. We also thank the laboratories and consortia that contribute to the public resources used in this manuscript. We are especially grateful to Rohit Sharma and the Mootha lab for devoting their 2020 lockdown time to enhance MitoCarta and creating the detailed gene-to-pathway annotations in MitoCarta3.0, which are fundamental to this work. This work was supported by NIH grants R01AG066828, R01AG086764, R35GM119793, the Wharton Fund, and Baszucki Group to M.P.

## SUPPLEMENTAL FIGURES

**Figure S1: Overview of the multi-tissue datasets and data processing.**

**A)** Mouse MitoCarta3.0 proteomics data. 3729 missing values were imputed as half of the minimal expression value of each protein, if the protein was found in at least one tissue. Protein data was used as is without added MitoPathway annotations.

**B)** Human protein atlas (HPA) tissue consensus dataset (version 21). This dataset encompasses one value per tissue, summarized across the HPA and GTEx transcriptomics datasets. Two genes are missing in the final dataset: RP11_469A15.2 (not found), and TSTD3 (missing values in several tissues). MitoPathway annotations were added for downstream analyses.

**C)** GTEx multi-tissue and -donor dataset (version 8). This dataset contains transcriptomics data from 54 tissues and 948 donors. Raw data was TMM normalized^73^ and 1133 MitoGenes were found. MitoPathway annotations were added for downstream analyses.

Abbreviations: NA, missing value; HPA, Human Protein Atlas, GTEx, Genotype-Tissue Expression project; TMM normalization, trimmed mean of M-values normalization.

**Figure S2: Fatty acid oxidation (FAO) specialization across the mammalian body**

**A)** Mouse MitoCarta proteomics ranked FAO scores quantified as a mean score of 42 proteins from MitoCarta by tissue. Fold changes per tissue relative to brain tissue average (4 tissues), indicating that anabolic tissues rank on average 10.2x higher in FAO scores than brain tissues.

**B)** Transcriptomics FAO scores in the HPA dataset showing a comparable signature in most tissues with skeletal muscle as exception. Liver ranks highest in FAO.

**C)** GTEx cortex vs liver mean difference across 226 liver and 255 cortex samples. Columns represent mean ± SEM. Welch two sample t-test, \*\*\**p*<0.001.

**D)** Individual FAO scores showing lowest and highest values in cortex and liver. The highest FAO score in the brain approaches the lowest FAO score in the liver. Fold changes indicate that liver shows a greater FAO variability than brain tissues.

**E)** Heatmap of FAD-dependent enzymes showing enrichment in liver tissues compared to cerebral cortex. Only PRODH is higher expressed in the brain. Gene expression was z-scored across liver and cortex samples.

**F)** Bivariate plot of FAD-dependent enzyme and complex I subunit scores showing that the liver mitotype is enriched for FAD-dependent enzymes relative to the brain.

**G)** Tight coupling of CII and FAO expression in the liver (spearman’s ρ = 0.0754, *p*< 0.001). The cortex does not show the same coupling (ρ 0.0925, ns).

**Figure S3: Comparison of normalization methods.**

**A)** Library sizes (i.e. total number of reads per sample) differ 5-13 fold in four human tissues: cerebral cortex, liver, skeletal muscle representing tissues with high mitochondrial content, and whole blood representing tissues with low mitochondrial content. Each column represents one sample, color coded by library size (small library size = yellow, large library size = dark blue).

**B)** Comparison of TPM (transcripts per million, not corrected) and TMM (trimmed mean of M-values, library size corrected) normalization methods using density blots of the four tissues indicated in A). For each tissue and sample, three pathway scores were calculated: Complex I (left panel), Fatty acid oxidation (FAO, middle panel), and Complex II (right panel). Both normalization methods show similar trends, though some tissue- and pathway-specific differences can be seen. TMM normalization is used for all analyses in this manuscript.

**Figure S4 Comparing human MitoPathways between tissues.**

**A)** Rank ordering of all MitoPathway scores (n=149 Level 3 pathways, MitoCarta3.0) by the expression difference between brain (averaged across 14 brain tissues) and liver. Pathways are color-coded by major pathway groups (Level 1 pathways). Pathways with the greatest difference between liver (bottom) and brain tissues (top) were annotated.

**B)** Urea cycle score vs Glycerol 3 phosphate (G3P) shuttle score across 55 human tissues, showing that liver mitochondria specialize in the Urea cycle while de-prioritizing the G3P shuttle.

**C)** BH4 synthesis vs serine metabolism score showing that brain mitochondria specialize in BH4 synthesis, while liver mitochondria prioritize serine metabolism.

**D)** MitoPathway radar chart of four tissues representing four major groups (brain, anabolic, contractile muscle, and digestive). For each tissue, MitoPathway scores were calculated, and the data was z-score-transformed within each pathway across the entire dataset. Top pathways for each of the four tissues were isolated (highest zscore across all tissues) and visualized. The boundaries of the radar chart are the highest (serine metabolism in liver, zscore = 7.3) and lowest (mt-mRNA modifications in small intestine, zscore = −0.8) values +/- 10%.

**E)** Bar chart of absolute MitoPathway scores from top pathways shown in D. Although the absolute scores show which tissue dominates in comparison to the other tissues, absolute scores between two pathways should not be directly compared, as they can be confounded by multiple transcripts resulting from higher mitochondrial content (see methods for details).

Abbreviations: G3P, Glycerol-3-phosphate; BH4, Tetrahydrobiopterin; metab., metabolism; SKM, skeletal muscle; CoQ, CoenzymeQ; VitA, Vitamin A; CI/II/III/IV, Complex I/II/III/IV; Anab., anabolic.

**Figure S5: Complex I (CI) ratios.**

**A)** CI / CII mean ratios (±SEM) across tissue groups. Welch’s t-test and Hedges’ g for effect size calculated between brain and all other groups.

**B)** CI / CIV mean ratios (±SEM). Student’s t-test and Hedges’ g for effect size calculated between brain and all other groups.

**C)** CI / CV mean rations (±SEM). Brunner-Munzel test and Hedges’ g for effect size calculated between brain and all other groups.

**D)** Ratios of CI vs all other pathways, each dot represents one pathway pair for one tissue. Some top pathways (i.e. where CI expression is greater) are highlighted.

**E)** Average of D (±SEM), indicating that brain and heart tissues prioritize CI 4.3x greater than other tissues. Abbreviations: CI/II/IV/V, Complex I/II/IV/V.

**Figure S6: mitoPPS calculations.**

**A)** MitoPathway ratio calculations across different mitospaces as visualized in Figure 3B.

**B)** High pathway1 (PW1) and low pathway2 (PW1) indicate priority shifts towards PW1, while low PW1 and high PW2 indicates deprioritization of PW1 and a priority shift towards PW2.

**C)** Estimating pathway prioritization from pathway ratios. For each tissue and pathway pair, a ratio is being calculated (left panel) and divided by the average ratio of the respective pathway pair across all samples (middle panel), resulting in corrected ratios (right panel).

**D)** Corrected ratio calculations are applied for all possible pathway combinations (here 148 for CI vs all others) and for all tissues (with cerebral cortex and liver as example). For each tissue and pathway, corrected ratios are averaged, resulting in Mitochondrial Pathway Priority Scores (mitoPPS).

**E)** Example CI mitoPPS across 55 human tissues indicating that brain and heart tissues show a high priority towards CI relative to all other tissues. Given that ratios are corrected (i.e. divided by a norm factor), mitoPPS scores are on the same scale for all pathways, with a score of 1 indicating average priority, a score of >1 higher priority, and <1 indicating lower priority. Abbreviations: PW, MitoPathway; CI, Complex I; CII, Complex II; FAO, Fatty acid oxidation;

**Figure S7: mitoPPS vs MitoPathway scores.**

**A)** Detailed view of the heatmap shown in Figure 5 with labeled MitoPathways.

**B)** Heatmap of MitoPathway z-scores with same order rows and columns as A), allowing to directly compare the two scoring methods. Tissues with higher average mitochondrial gene expression (shown rank-ordered in **C**) usually have the highest MitoPathway zscore, which is not the case for mitoPPS.

**C)** Spearman’s correlation between mitoPPS and MitoPathway scores across all pathways, only mtDNA pathways (i.e. pathways composed of genes encoded in the mtDNA) and non-mtDNA pathways (i.e. pathways not composed of mtDNA genes). The strongest correlation is within mtDNA pathways, followed by non-mtDNA pathways. All pathways together correlate weakly, given that MitoPathway scoring does not correct for the transcript-enrichment in mtDNA pathways, which in contrast mitoPPS does. \*\*\**p*<0.001.

**Figure S8: Specialization across the human body.**

Detailed barplots of the mitoPPS shown in Figure 5F across all 55 tissues. The data is log10-transformed, and a log10(mitoPPS) of 0 means average priority, >0 indicates higher priority, and <0 lower priority.

**Figure S9: Cellular lifespan study mitoPPS**

**A)** Processing of the Cellular lifespan study RNAseq data. One gene was missing in the dataset (RP11_469A15.2).

**B)** Detailed view of the heatmap shown in Figure 6 with all MitoPathways as rows and conditions/samples as columns, sorted by condition and ascending passage (see top annotation).

**C)** PCA of multi-tissue and fibroblast data with controls, treatments, and the genetic OxPhos defect caused by a mutation in the SURF1 gene, color-coded by passage. Given that the passage-dependent shift (shown in Fig6C) is also present in the treated fibroblasts, the data was averaged across all passages for each condition for downstream analyses.

**D)** PCA of multi-tissue and averaged fibroblast data across experimental conditions, showing some treatment group-specific clustering. For three conditions (2DG, BHB, Galactose), data from only 2 cell lines was available. All other treatments contain data from three cell lines, either from healthy controls or with a SURF1 mutation.

**E)** 3D plot of the PCA shown in D with PC3. Oligomycin and Oligomycin + Dexamethasone - treated fibroblasts cluster apart from other treatments along PC3.

**Figure S10: Comparison of tissue and fibroblast mitoPPS**

**A)** Radar charts of complex I mitoPPS for a subset of tissues (left, grey) and fibroblasts (right, blue). The scales of both radar charts are the same, scaled by the maximum and minimum value of both datasets (100% = maximum value, from cerebral cortex; 0% = minimum value, from bone marrow). Light grey indicates the average across the entire dataset for tissues and fibroblasts respectively (∼1, i.e. equal priority, <1 deprioritization, >1 prioritization). Labels contain percent changes relative to average for each dataset.

**B)** Radar charts of Fatty acid oxidation,

**C)** Radar chart of mtDNA repair in tissues (left panel) and scatter plot of mtDNA repair in control fibroblasts, split by cell line (hFB12, hFB13, and hFB14). Percent changes in the scatterplot are calculated relative to the earliest passage.

**Figure S11: Dynamic range of tissue and fibroblast mitoPPS and correlation with bioenergetics.**

**A)** Detailed barplot of dynamic mitoPPS range shown in Figure 6H with pathways as rownames.

**B)** Bivariate plot of Catechol metabolism vs Phospholipid metabolism mitoPPS in tissues and fibroblasts. While tissues do not prioritize the two pathways, fibroblasts treated with the synthetic glucocorticoid Dexamethasone or mitochondrial OxPhos defect inducers Oligomycin and MitoNUITs shift their priority to the catechol or phospholipid metabolism pathway, respectively.

**C)** Bivariate plot of Iron homeostasis vs Cholesterol-associated mitoPPS showing that Oligomycin-treated cells prioritize Iron homeostasis, while cells from the forced OxPhos and contact inhibition groups prioritize both pathways alike, relative to tissues and all other treatments.

**D)** Correlation of baseline OxPhos-linked ATP production rate (ATPox) with mitochondrial central dogma mitoPPS across treatment conditions (averaged per treatment and cell line, ρ=-0.76, p.adj<0.001).

**E)** Correlation of ATPox with cholesterol-associated pathway (ρ=0.77, p.adj<0.001).

**F)** Heatmap showing Spearman’s correlations between mitoPPS values for each pathway and multiple Seahorse-derived bioenergetic parameters across all treatment conditions. \**p*<0.05, \*\**p*<0.01\*\*\**p*<0.001. p-values were adjusted using the Benjamini-Hochberg (BH) correction for multiple testing.

**Figure S12: Detailed presentation of the data shown in Figure 5**

**A)** The same heatmap shown in Figure 5E, annotated with pathway names for clarity.

B) Principal component analysis plot similar to 5D, but including all ROSMAP individuals rather than group averages. Corresponding MitoBrainMap samples are highlighted in circles.

**Figure S13. Cell-type annotation and validation of germ cell markers in the adult human testis single-cell RNA-seq dataset ^55^.**

**A)** UMAP visualization of the adult human testis scRNA-seq dataset from Guo *et al.* showing the major annotated cell populations, including spermatogonia, spermatocytes, spermatids, Sertoli, Leydig, peritubular, endothelial, and macrophage cells.

**B)** Seurat clusters of the same dataset, used for cell type annotations as reported in Guo et al.

**C)** Cell state assignments based on cluster identity, highlighting the continuous differentiation trajectory from spermatogonia to late spermatids.

**D)** Expression patterns of canonical germ cell markers across the UMAP embedding confirm correct cell state annotation. Early germ cell markers (*UTF1*, *FGFR3*, *ID4*, *KIT*) are enriched in spermatogonia; meiotic markers (*DMRT1*, *DMRTB1*, *SYCP3*, *SPO11*, *MLH3*) peak in spermatocytes; and post-meiotic markers (*ZPBP*, *TNP1*, *PRM2*) are expressed in spermatids, consistent with established developmental staging.

**Figure S14. Mitochondrial pathway dynamics along spermatogenesis.**

**A)** Expanded view of the heatmap shown in Figure 6E with all pathway annotations shown.

**Figure S15. Mitochondrial pathway dynamics along spermatogenesis.**

**A)** Expanded view of top-prioritized MitoPathways showing dynamic prioritization across early, mid, and late stages of spermatogenesis. Top-pathways are grouped by spermatogenesis phase (early, mid, late).

**B-E)** Representative mitoPPS trajectories modeled by generalized additive models. Curves show smoothed mitoPPS trajectories (mean +/- s.e.) along pseudotime, with vertical lines marking cell-state transitions.

## METHODS

### Lead contact

All code and data-related questions can be addressed to the lead contact Anna Sophia Monzel

### Data and Code Availability

All datasets used in this study are publicly available and can be accessed on the respective domain (see Data sources). The code is written in R version 4.5.1 (2025-06-13), and can be accessed through GitHub, together with the package versions (https://github.com/annamonzel/mitotyping ).

### Data sources

This study uses multiple publicly available datasets. The human and mouse MitoCarta3.0 datasets were obtained from the Mootha lab at the Broad Institute^20^.

Tissue-level transcriptomic data were obtained from the Human Protein Atlas (HPA) tissue consensus RNA dataset, which integrates RNA-seq data from both HPA and GTEx. For each gene, the normalized transcript expression (“nTPM”) represents the maximum nTPM value across the two resources. The dataset provides one expression value per gene and tissue and was downloaded from HPA (https://v21.proteinatlas.org/about/download at version 21.0). Notably, newer release (v21.1+) do not include the olfactory bulb. However, all analyses were repeated excluding this tissue, and results were unchanged. This information can also be found in the associated code.

GTEx liver and cerebral cortex data used in Figure 2H-K and Supplemental Figures S2 C-G and S3 were obtained from the GTEx consortium (948 donors, 17382 samples, of which 226 were liver donors, and 255 cortex donors; https://www.gtexportal.org).

The cellular lifespan RNA-seq dataset was obtained from previous work^44,45,47^ via Gene Expression Omnibus (GSE179848). The normalized RNAseq data and Seahorse bioenergetics data are also accessible through the Lifespan Study Shinyapp (https://columbia-picard.shinyapps.io/shinyapp-Lifespan_Study/), though RNA-seq normalization methods differ from those used here.

Single-nucleus RNA-seq data from the human MitoBrainMap ^49^ were accessed at http://humanmitobrainmap.bcblab.com/.

Post-mortem single-nucleus RNA-seq data from dorsolateral prefrontal cortex of donors in the Religious Orders Study and the Memory and Aging Project (ROSMAP) ^50^ were accessed via the AD Knowledge Portal https://adknowledgeportal.synapse.org/.

Single-cell RNA-seq data from the human testis^55^ was accessed through the HPA Single cell type data portal (https://www.proteinatlas.org/humanproteome/single+cell/single+cell+type/).

### Data pre-processing

#### Mouse MitoCarta3.0

The mouse MitoCarta3.0 proteomics dataset of 14 tissues was used as is (total peak intensity). The dataset misses many proteins that are either tissue specific, show different temporal expression patterns, are part of the mitochondrial outer membrane^20^, or we reasoned they are below the detection limit. Since missing values are problematic for hierarchical clustering and principal component analysis, we imputed some, yet not all, missing values. First, we removed proteins that were not detected in any of the 14 tissues (163 proteins). For the remaining proteins, we reversed the log10 transformed data, and imputed NAs as half of the minimal value of each gene to get a low value that is non-zero. In total, 3729 (27%) missing values were imputed, resulting in a total of 977 proteins in 14 tissues. We next assigned groups to tissues with functional similarities (**Supplemental table 1**).

#### Human MitoCarta3.0

The human MitoCarta3.0 dataset served as both, a database of mitochondrial genes (both nuclearDNA- and mitochondrialDNA-endcoded, 1136 genes), as well as a resource for gene-to-pathway annotations (Supplemental File 1). Some genes play a role in multiple pathways, and the amount of genes within each pathway differ (see Supplemental table 2). Pathways exist on three levels that we outline here with OxPhos as example: the first level pathway “OxPhos”, contains multiple second level pathways (“Complex I”, “Complex II”,…), and multiple third level pathways (“Complex I subunits”, “Complex I assembly factors”). The latter two usually have no shared genes, but level 2 pathways contain all level 3 genes, and level1 pathways contain genes of levels 2 and 3. Thus, some pathways can show similar expression patterns due to shared genes.

#### Human Protein Atlas multi-tissue dataset

The Human Protein Atlas consensus tissue transcript expression dataset, based on transcriptomics data from HPA and GTEx^21,23,74^, was accessed at version 21.0 (2021.11.18) with ENSEMBL version 103.38 The data format was unchanged (nTPM = normalized transcripts per million, TMM normalization and 1134 mitochondrial genes in 55 human tissues were found. One gene was missing (RP11_469A15.2) and one gene (TSTD3) was excluded due to missing values in multiple tissues. Same as for the mouse proteomics dataset, we assigned groups to tissues with functional similarities as shown in Supplemental table 1. Any modification to the nTPM data (zscore-transform, log10 transform) is described in the respective figure legend. Given that the HPA dataset contains *one* value per tissue (summarized across two datasets from multiple donors), statistical power is limited. Multi-donor datasets such as the Genotype-Tissue Expression (GTEx) should be explored in future studies for robustness and to explore inter-individual variability. Another limitation of this (bulk transcriptomics) dataset is that the data is composed of cell type mixtures, likely with one dominating cell type (for example cardiomyocytes in the heart). As a result, such tissue-based analyses may underestimate the true magnitude of differences between pure mitotypes. Resources such as the Human Cell Atlas^75^ can further our understanding of cell-type level mitotypes in human tissues.

Before conducting analyses, tissues were assigned to tissue groups based on functional similarity to our best knowledge, and this classification was maintained throughout the study. When classification was unclear, tissues were assigned to the “other” category. For example, the tongue was placed in the “other” category due to uncertainty about its specific sampling location. Although it clustered with “contractile muscle”, we retained our original classification throughout the study.

#### GTEx multi-individual dataset

We used GTEx version 8 (2017-06-05)^23,71^ raw count data (gene_reads) and applied TMM normalization using the edgeR package^73^ . Although the data can be accessed as transcripts per million (TPM), we reasoned that varying library sizes (i.e. the total number of mapped gene reads) can lead to over- or underestimation of MitoPathway scores. We demonstrate the differences in library sizes using four distinct tissues (brain – cortex, liver, skeletal muscle, and whole blood, **Figure S3**). Library sizes within tissues can vary by 13-fold, and whole blood library sizes are generally lower compared to the other three tissues. This is most likely due to lower amounts of mtDNA-encoded transcripts and fewer copies thereof, further confirming the necessity to correct for it^76,77^. We next compared the effect of normalization using histograms for each of the four tissues and MitoPathway scores of the three pathways studied in GTEx (Complex I, Fatty acid oxidation, and Complex II). While the directionality between the uncorrected (TPM) and library-corrected (TMM) pathway scores is largely similar, the data distribution within tissue groups varies between the two methods. For example, Complex II expression is low in brain and whole blood, compared to liver and muscle. While the whole blood Complex II TPM score is skewed towards lower expression, TMM normalization corrects this towards a normal distribution, while brain and whole blood still remain lowest.

After TMM normalization, we pulled all mitochondrial genes from the GTEx dataset using ENSEMBL mapping between MitoCarta and GTEx (see code for details). In total, we identified 1133 genes in 54 tissues and 928 donors. The three missing genes were SOC2, CMC4, and ATP5MF-PTCD1. For GTEx mitotyping (Figure 2 F-I) we used only data from the cerebral cortex (n=255 donors) and liver (n=226 donors), among which 79 cortex and liver samples were from the same donor.

#### Cellular lifespan study

The unprocessed cellular lifespan dataset (GSE179848) was imported using txi import (length-scaled TPM), and further TMM normalized using the NOISeq package. The resulting nTPM values are the TMM normalized protein-coding transcripts per million data. 1135 mitochondrial genes were found (RP11_469A15.2 missing). We noticed that mtDNA-encoded genes such as MT-ND1 and MT-ND2 were missing the “MT-“ suffix, which was added manually. In addition, gene C12orf10 was renamed to its synonym MYG1 for mapping. One sample (RNAseq_Sample152, part of Figure 6H) was excluded due to infinity values in MitoPathway scores (see code for details). During the course of the experiments, we noticed that one (apparently healthy) control line used (HC3, Coriell AG01439) was likely isolated from a diseased newborn, since the infant had died of unknown cause 4 days after birth^45^. In total, 399 samples from 7 cell lines and 15 conditions (genetic differences or treatments or both) were mitotyped (see **Supplemental File 4**).

#### MitoBrainMap

The clean and downsampled snRNAseq seurat object from the human MitoBrainMap was loaded, and the white matter sample (“voxel 4”), as well as vascular and leptomeningeal cells (VLMCs) were excluded to match the ROSMAP snRNAseq dataset, which is grey matter only and lacks VLMCs. A new UMAP object was generated using the filtered dataset (Figure 5A). Gene expression matrices were pseudobulked per cell type x brain region pair using Seurat’s AverageExpression() function. MitoPathway scores were calculated as described in detail in ^49^. In brief, 1118 mitochondrial genes were found, and MitoPathway scores were calculated for each cell type / original voxel using MitoCarta3.0 annotations.

### ROSMAP

Single-nuclei RNA seq data was pseudobulked using aggregated single-nucleus expression for each cell type and donor, and the resulting log2-normalized counts per million were used to calculate MitoPathway scores. A total of 1090 mitochondrial genes were found.

#### Human testis cell atlas

Single-cell RNAseq data from human testis ^55,74^ were analyzed using Seurat v5.3.0. Cell-level read counts and metadata were merged with cluster annotations from the original study. Ensembl gene identifiers were mapped to gene symbols, and duplicated mappings were collapsed by summing counts. Genes expressed in <0.1% of cells and with <10 total UMI counts across all cells were removed to filter low-abundance transcripts.

*Normalization and dimensionality reduction:* Gene expression values were normalized using NormalizeData() and highly variable genes were selected using FindVariableFeatures() (2,000 features). Data were scaled and principal component analysis (PCA) was performed. UMAP embeddings supplied with the dataset were incorporated directly and stored as a dimensional reduction object in Seurat. PCA-derived UMAP embeddings (30 PCs, min.dist = 0.3, n.neighbors = 30) were computed for refined clustering shown in the main figures.

*Cell-type and subtype annotation:* Cells were visualized and clustered by annotated high-level cell classes (**Figure S13A**). Germ-cell subsets (spermatogonia, spermatocytes, early spermatids, late spermatids) were retained (**Figure S13B**), and clusters were re-assigned to germ-cell subtypes according to published annotations from Guo et al^55^ (**Figure S13C**). Annotation accuracy was verified by inspecting expression patterns of canonical markers (e.g., UTF1, ID4, FGFR3 for spermatogonial stem cells; KIT, DMRT1, SYCP3 for spermatocytes; TNP1, PRM2 for spermatids, see **Figure S13D**).

*Trajectory inference:* To infer a developmental trajectory through spermatogenesis, the processed Seurat object was converted to a SingleCellExperiment object and pseudotime inference was performed using **Slingshot** v2.16.0 with UMAP embeddings as input. Lineage-specific pseudotime values were extracted and stored for downstream analyses and visualization. Cells not assigned a pseudotime value were excluded for downstream analyses.

*Cell cycle scoring:* Cell cycle phase scores (G2/M, S phase) were calculated using Seurat’s CellCycleScoring() function with canonical gene sets and added to the metadata.

*Pseudotime binning:* Cells were ordered by pseudotime, excluding unassigned cells. Cells were grouped into consecutive bins of 10 cells along pseudotime. For each bin, mean pseudotime and the dominant germ-cell subtype were computed. A cell-to-bin dataframe was exported and used for downstream grouping.

*Gene sets:* MitoPathway gene sets were curated from MitoCarta3.0. To maximize gene recovery, missing genes were recovered via i) the MitoCarta synoym table and ii) the checkGeneSymbols() function from the HGNChelper package. All matches were manually verified. Two non-mitochondrial pathways were included:

- Glycolysis: HK1, HK2, HK3, GCK, GPI, PFKL, PFKM, PFKP, ALDOA, ALDOB, ALDOC, TPI1, GAPDH, PGK1, PGAM1, PGAM2, ENO1, ENO2, ENO3, PKM, PKLR
- Pentose phosphate pathway: G6PD, PGLS, PGD, RPIA, RPE, TKT, TALDO1 Final gene lists were saved for downstream gene set enrichment analyses.

*MitoPathway scores:* Mitochondrial and glycolysis/ pentose phosphate pathway gene sets were passed to the Seurat AddModuleScore() function to compute pathway scores per cell based on the average expression of pathway genes relative to control gene sets. Since strictly positive scores are required for ratio-based metrics, pathway scores were shifted by subtracting the global minimum and adding a small constant (1×10-6) without altering relative variation. Resulting module scores were appended to the Seurat metadata and averaged across the previously assigned pseudotime bins to obtain bin-level mean pathway scores.

### Quantification and Statistical analyses

#### Mouse FAO score

A fatty acid oxidation score was calculated using expression data from the following genes: Acaa1a, Acaa2, Acacb, Acad10, Acad11, Acad12, Acadl, Acadm, Acads, Acadsb, Acadvl, Acat1, Acot11, Acsf2, Acsl1, Acsl6, Acsm1, Acsm2, Acsm3, Acsm4, Acsm5, Acss1, Amacr, Cpt1a, Cpt1b, Cpt1c, Cpt2, Crat, Crot, Decr1, Echs1, Eci1, Eci2, Etfb, Etfdh, Hadh, Hadha, Hadhb, Hsd17b10, Mcee, Mmut, Pcca, Pccb, Slc25a20. *A* score was calculated using the average normalized expression of the aforementioned genes.

#### Mitopathway scores

As mentioned above, gene-to-pathway annotations were extracted from MitoCarta3.0. To calculate MitoPathway scores from any normalized expression data (HPA, GTEx, Cellular lifespan study), expression values of all genes within a given pathway were averaged. Pathways that contain mtDNA-encoded genes always rank highest due to multiple copies of the same gene. Hence, MitoPathway scores should be expressed in *relative* terms (e.g. relative to all other tissues, e.g. which tissue ranks highest in MitoPathway Complex I). Importantly, the scoring requires that most mitochondrial genes associated with a given pathway are represented in the dataset. Insufficient gene coverage can affect the reliability and biological relevance of the calculated scores, potentially leading to inaccurate interpretations.

Any modification to the data (zscore, log10 transform, log2 fold changes, ratios) is specified in figure legends. Typically, data was zscore-transformed within pathways, log2 fold changes were calculated between two tissues and ratios were calculated between two pathways for each tissue.

#### mitoPPS

Given that MitoPathway scores differ by multiple orders of magnitude (e.g. complex I scores spans 5 orders of magnitude, while fatty acid oxidation scores span 2 orders of magnitude from lowest to highest tissue), absolute MitoPathway scores are not directly comparable. In addition, tissues with a naturally greater mitochondrial content rank highest in most MitoPathways. To address this scaling issue, we used a normalized ratio-based approach that determines how much a sample prioritizes a MitoPathway over all other MitoPathways and samples. We eliminated the effect of overall mitochondrial gene expression level by dividing individual MitoPathway score ratios by the average ratios across all samples of a defined reference population for each respective pathway pair. Thus, for each sample of interest S_i_ and pathway of interest P_i_ the respective mitoPPS is calculated as follows:

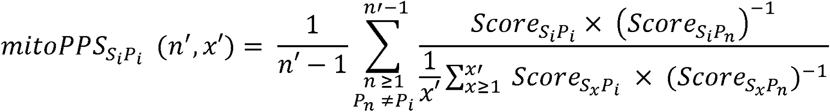

Where

S_i_= Sample of interest

P_i_= Pathway of interest

Score = MitoPathway score

n’ = Total number of Pathways

x’ = Total number of Samples

P_n_ = n^th^ pathway in set of pathways P excluding PI

S_x_ = x^th^ sample in a dataset

#### Dataset-specific implementation

*Human Protein Atlas:* Pairwise pathway ratios for each tissue were normalized to the mean ratio across all tissues to remove tissue-driven differences in global mitochondrial abundance.

*Cellular lifespan study:* The same framework was applied, except ratios were normalized to the mean ratio of control (healthy, untreated) samples to quantify treatment- and aging-associated shifts in pathway prioritization. For Figure 6B-G we used control samples from the longest culture from study part 2^44^, but included all data in the heatmap in Figure 4H.

*MitoBrainMap:* mitoPPS was computed within brain regions, with pathway ratios normalized to region-specific averages. This preserved cell-type–specific prioritization rather than regional metabolic differences (see also ^49^)

*ROSMAP:* mitoPPS was computed per donor and cell type. For each donor–cell type pair, pairwise pathway ratios were calculated and normalized to the mean ratio across all donors for each pathway pair. This approach preserves inter-individual mitoPPS variation while controlling for overall pathway abundance differences.

*Spermatogenesis:* Pairwise ratios were computed per bin and normalized to the global mean ratio to quantify dynamic pathway prioritization along germ-cell development.

#### Spermatogenesis scGSEA scores

To verify pathway activity independent of mitoPPS, single-cell gene set enrichment analysis (scGSEA) was performed in R v4.3.0 (on Ubuntu Linux) using the gficf package (v2.0.0)^63^. Raw UMI counts were normalized using gficf(), and enrichment scores were computed with runScGSEA() using 151 random seeds (equal to the number of pathways) to account for algorithmic stochasticity. Gene-to-pathway annotations from MitoCarta3.0 and manually curated pathways (glycolysis and pentose phosphate pathway) were used as input, with a false discovery rate (FDR) threshold of 0.05. Within each run, pathway scores were z-scaled across all cells. Pathways enriched in >10% of runs were retained, and final pathway scores were calculated as the mean z-score across runs for each cell.

The resulting cell-by-pathway matrix was imported into a Seurat object alongside the original UMAP embeddings for visualization. Scores were visualized using FeaturePlot().

#### Spermatogenesis mtDNA vs nDNA dynamics

Log-normalized single-cell expression values were extracted from the Seurat object and merged with pseudotime metadata. Mitochondrial genes were classified as either mtDNA-encoded (genes prefixed with MT-) or nuclear-encoded mitochondrial genes (nDNA, all others). Mean expression per genome group was computed for each cell, then averaged across pseudotime bins. Bin-wise trajectories were smoothed (rolling window, k = 80 bins) and z-scaled to allow direct comparison of dynamic patterns. Germ-cell stage transitions were marked based on dominant cell-type labels along pseudotime.

#### Spermatogenesis top pathways

To identify mitochondrial pathways with dynamic prioritization during spermatogenesis, bins with mitoPPS > 1.1 were considered activated (i.e. 10% higher compared to average priority). Pathways were retained if they showed activation in ≥ 10 bins and a dynamic range (max–min) ≥ 0.4 (i.e. 40% change along spermatogenesis).

Retained pathways were classified as Early, Mid, or Late by determining the developmental phase (based on dominant cell-type labels along pseudotime) in which > 30% of activated bins occurred.

#### Hierarchical clustering + PCA

All heatmaps were generated using the ComplexHeatmap package^78,79^. Data transformations (z-score, log transform) can be found in the figure legends. For hierarchical clustering, the Euclidean distance was calculated, and cluster analysis was performed using the Ward’s D2 hierarchical Agglomerative Clustering Method^72^. If a different method (such as k-mean clustering) was used, the information can be found in the figure legends.

Principal component analysis was conducted using the base R prcomp function. Loadings can be found in the supplemental information. 3D plots of the principal components were generated using the rgl package and the plot3d function.

#### Radar charts

Radar charts were generated from z-score transformed (MitoPathwayScores) or mitoPPS data using the fmsb package. The boundaries of the radar chart are scaled within each dataset, with 100% representing the highest value (+10%) and 0% representing the lowest value (−10%).

#### Statistical tests

All statistical tests relevant to the figures can be found in the figure legends. Statistics not reported in the figures (but reported in the text) can be found in the associated code on GitHub. All correlations in this manuscript were performed using Spearman’s rank-order correlation with the cor.test function (stats package). P-values were adjusted using the Benjamini-Hochberg procedure. Effect size was calculated using Cohen’s D with Hedges’ g correction using the cohen.d function from the effsize package (hedges.correction=TRUE). When two groups were directly compared, first a Shapiro-Wilk test (stats package) was used to test for normal distribution, and a Fliegner-Killeen test (stats package) was used to test for equal variance. If the data was normally distributed and variances were homogeneous, a Student’s t-Test (stats package) was performed. If the assumption of equal variance was not met in a normally distributed set of data, a Welch’s t-Test was performed. Non-parametric tests were used when the data was not normally distributed. For homogeneous variance a Wilcoxon Rank Sum test (Mann Whitney, stats package) was performed. If the variance was not homogeneous, a Brunner-Munzel test (brunnermunzel package) was used.

### Additional resources

The code for each figure, including all statistical analyses can be found on GitHub (https://github.com/annamonzel/mitotyping)

