## Supplemental figures for "Mitotyping – An integrative framework to quantify mitochondrial specialization and plasticity"

#### SUPPL FIGURE S1

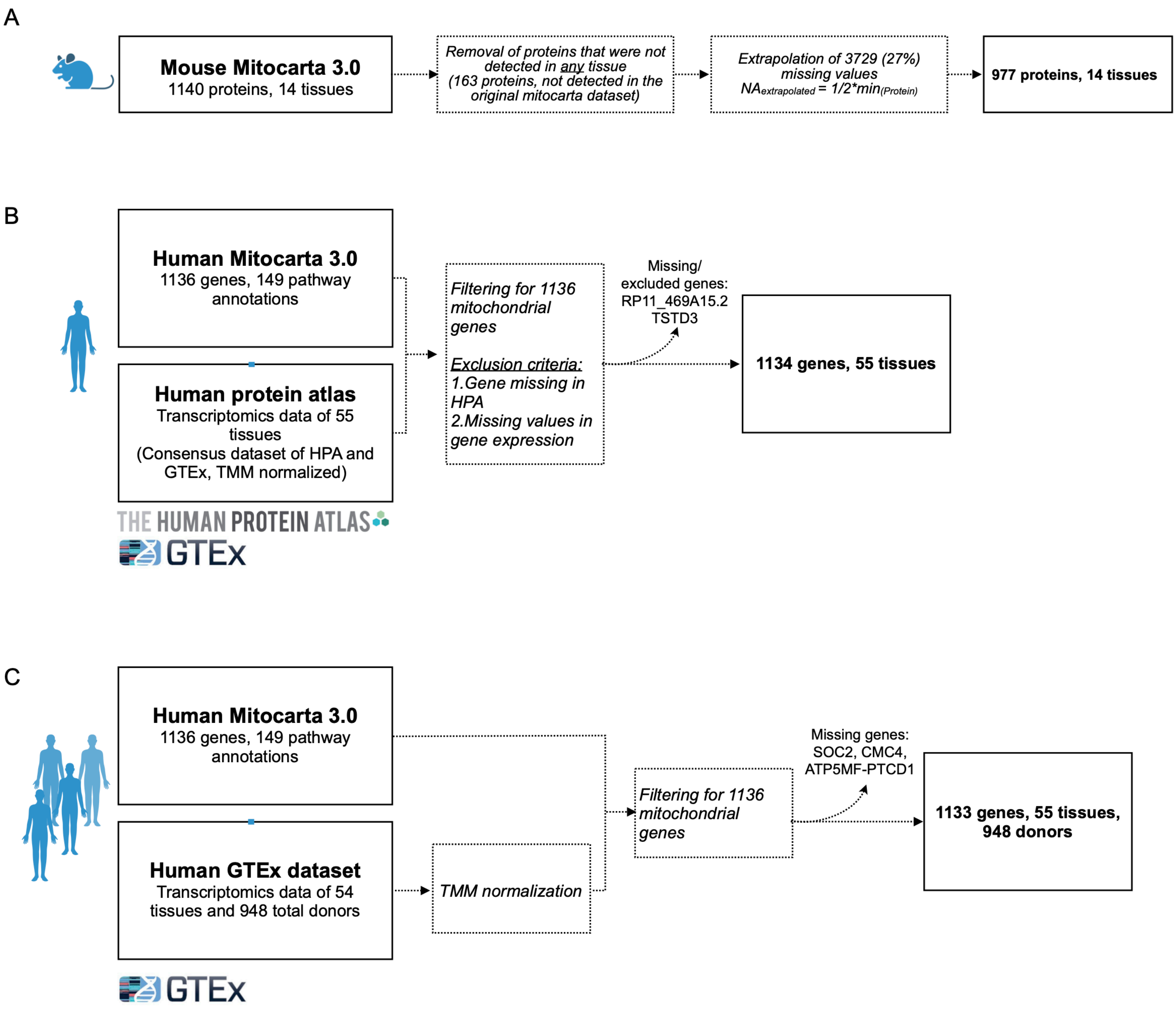

### SUPPL FIGURE S2

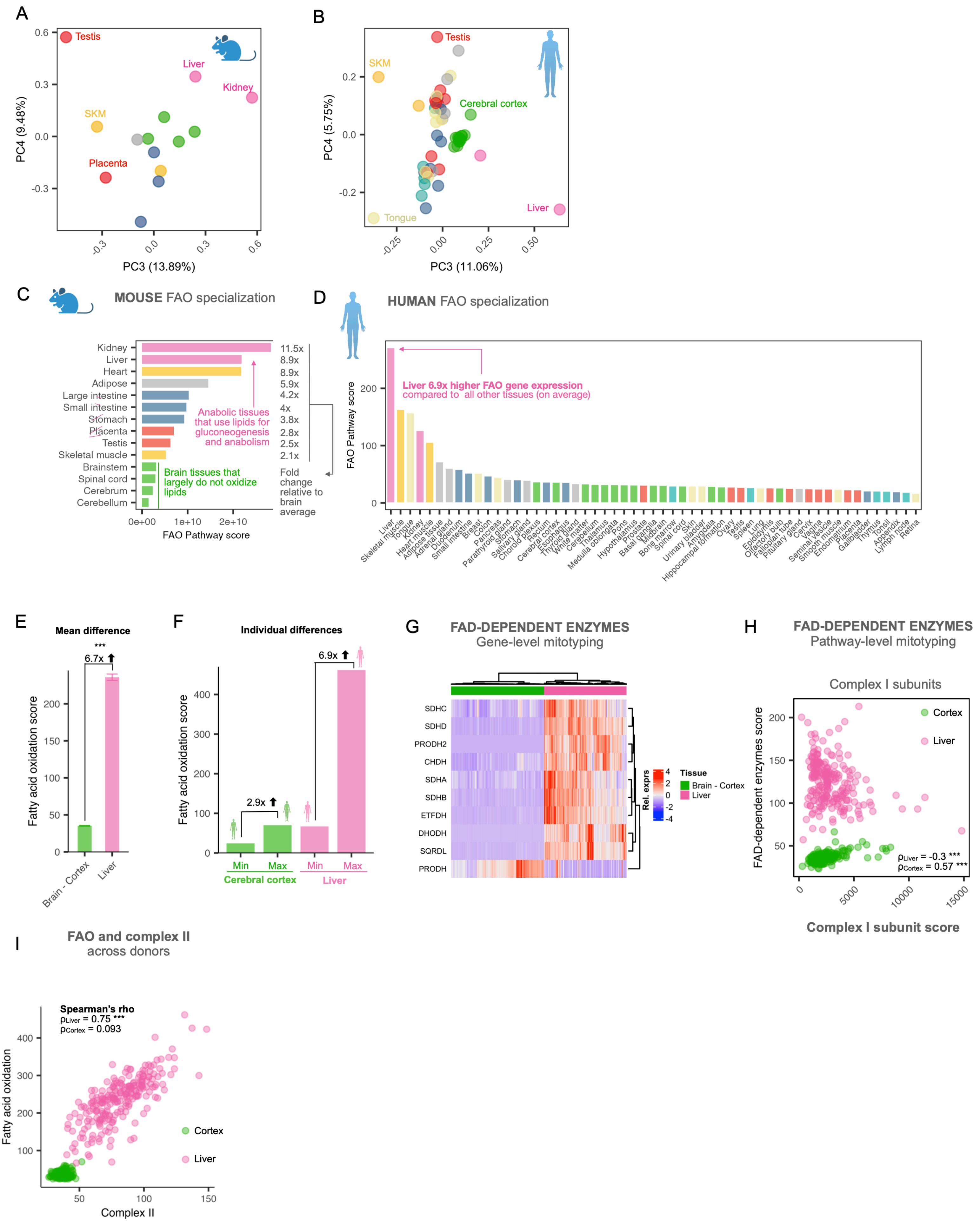

#### SUPPL FIGURE S3

A LIBRARY SIZE PER TISSUE FROM GTEx

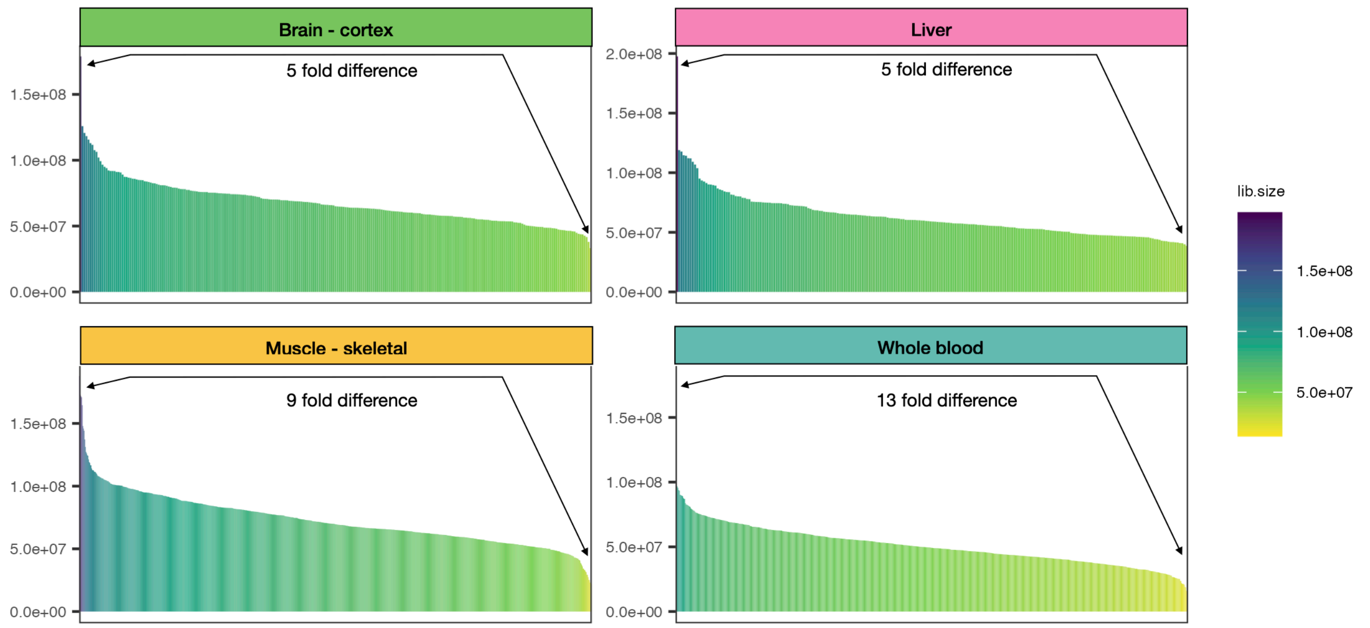


B COMPARISON OF NORMALIZATION METHODS

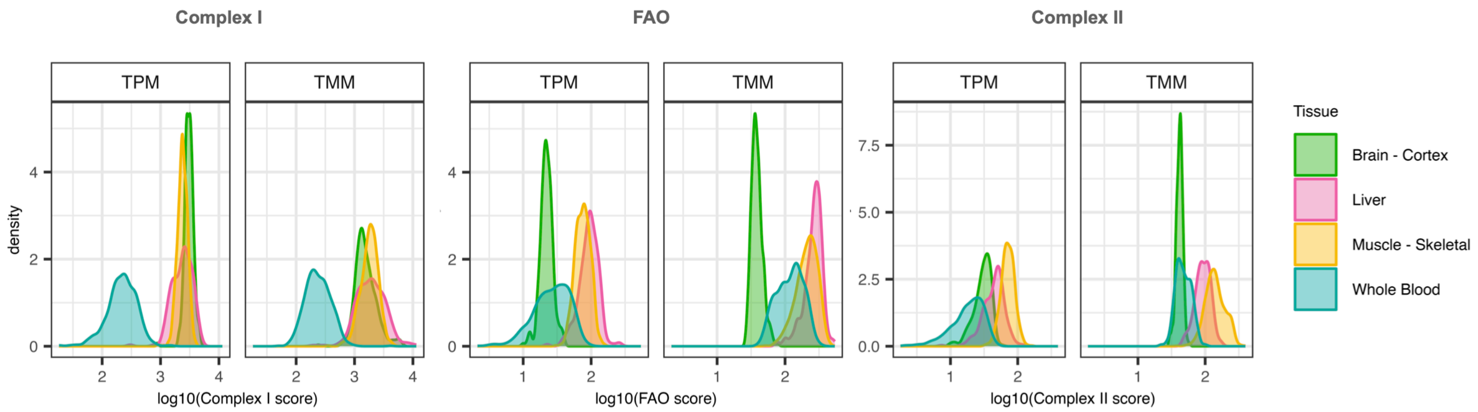

### SUPPL FIGURE S4

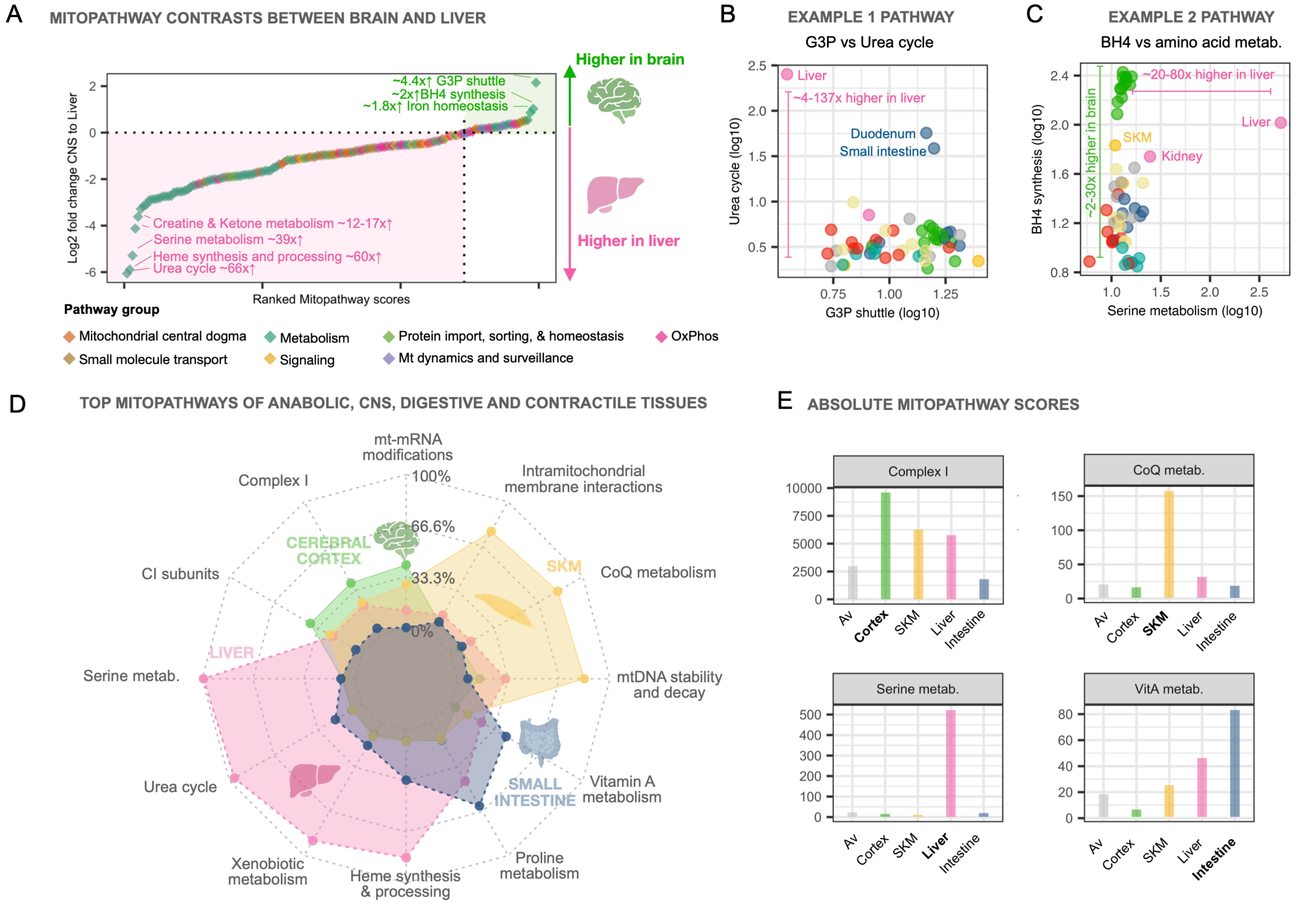

#### SUPPL FIGURE S5

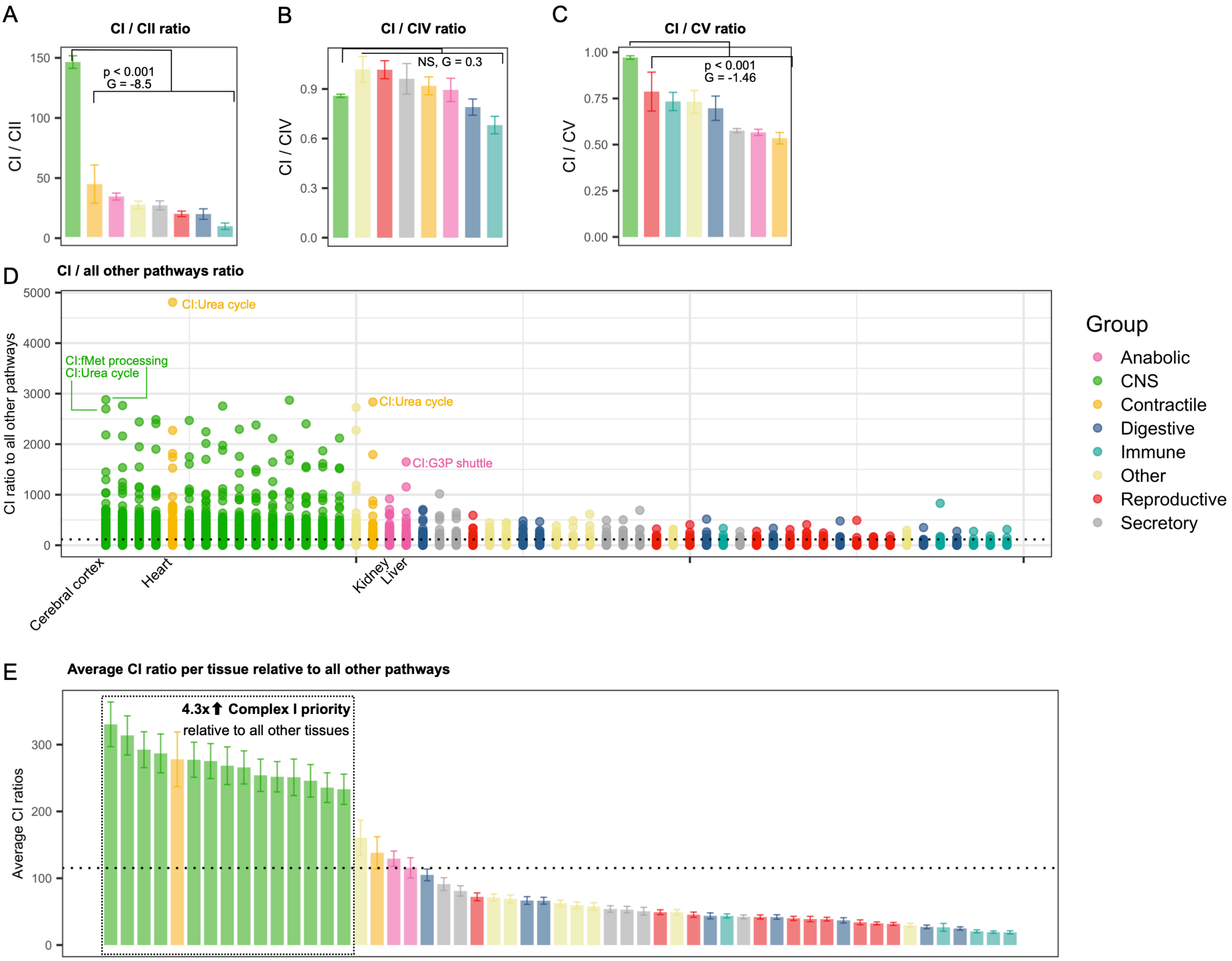

#### SUPPL FIGURE S6

A PATHWAY RATIOS

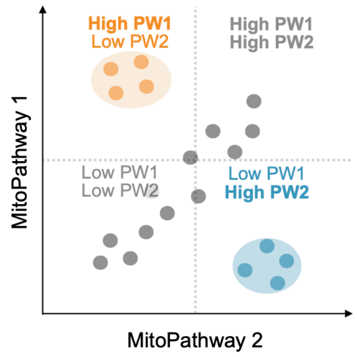


B PATHWAY PRIORITIZATION

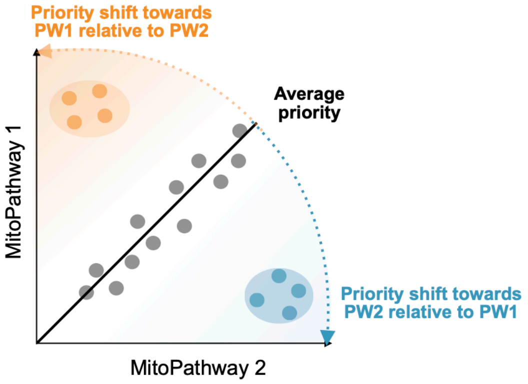


C ESTIMATING PATHWAY PRIORITIZATION FROM CORRECTED PATHWAY RATIOS

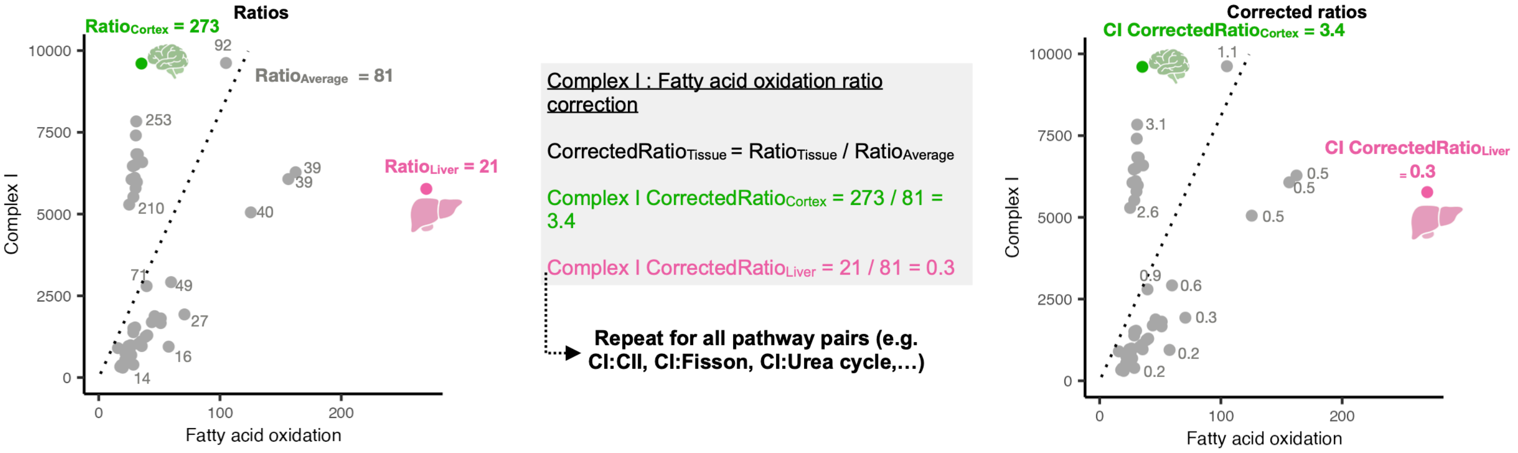


D mitoPPS: AVERAGED CORRECTED PATHWAY RATIOS ACROSS 148 PATHWAY PAIRS

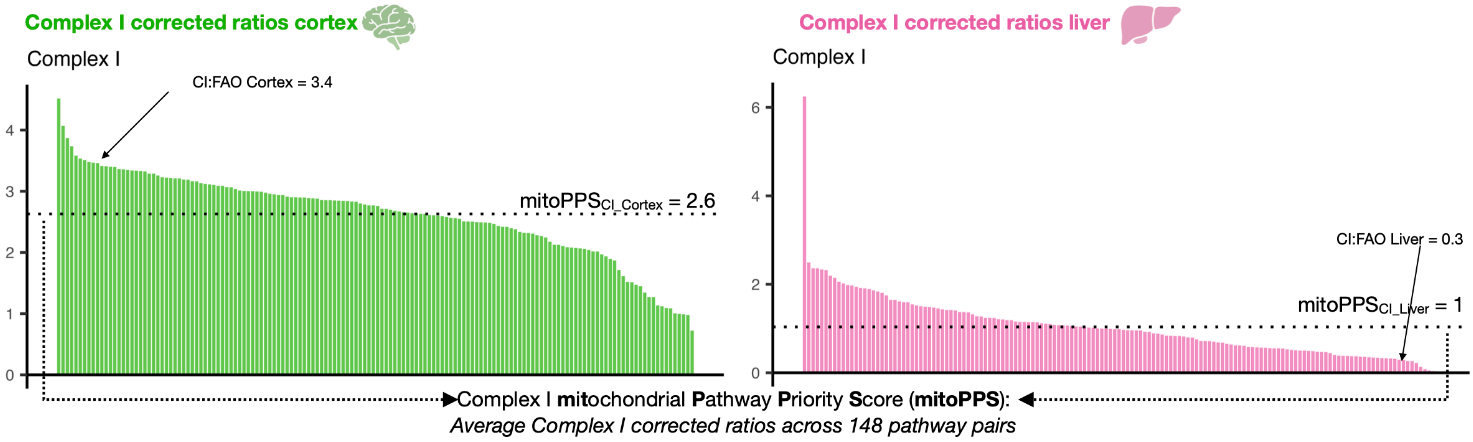


E COMPLEX I mitoPPS ACROSS 55 HUMAN TISSUES

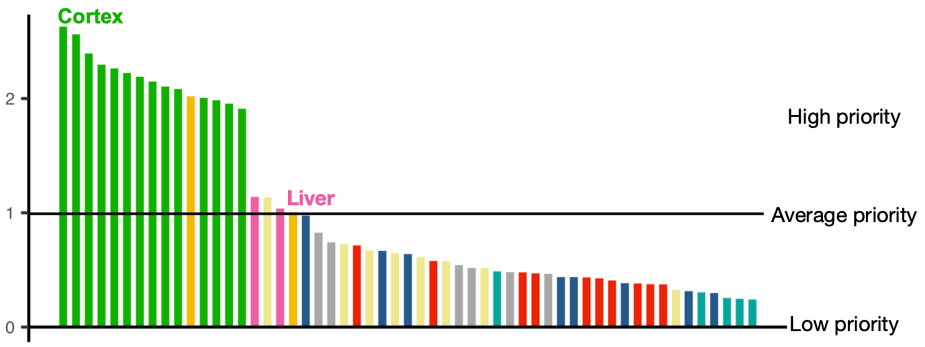

#### SUPPL FIGURE S7

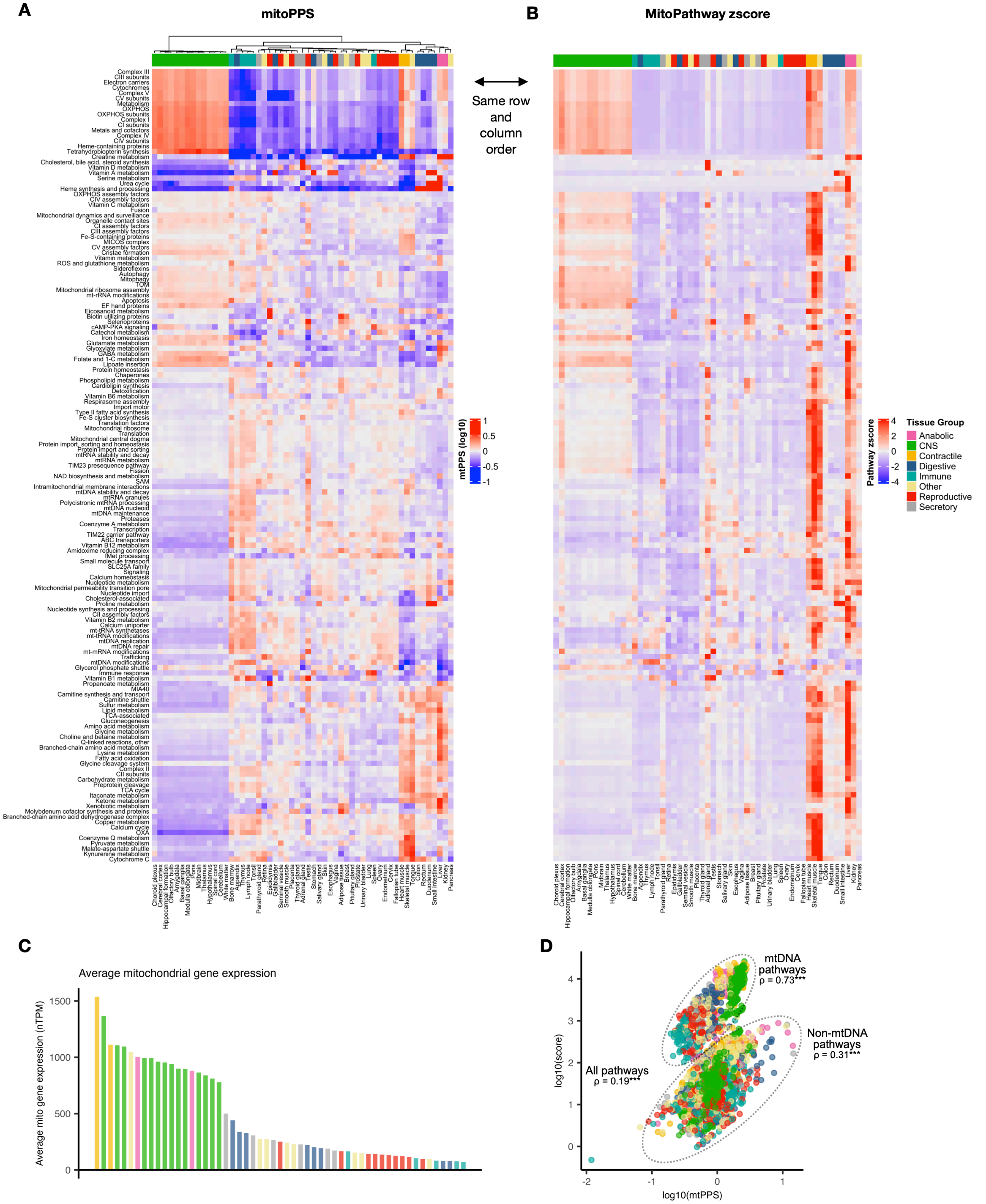

**SUPPL FIGURE S8**

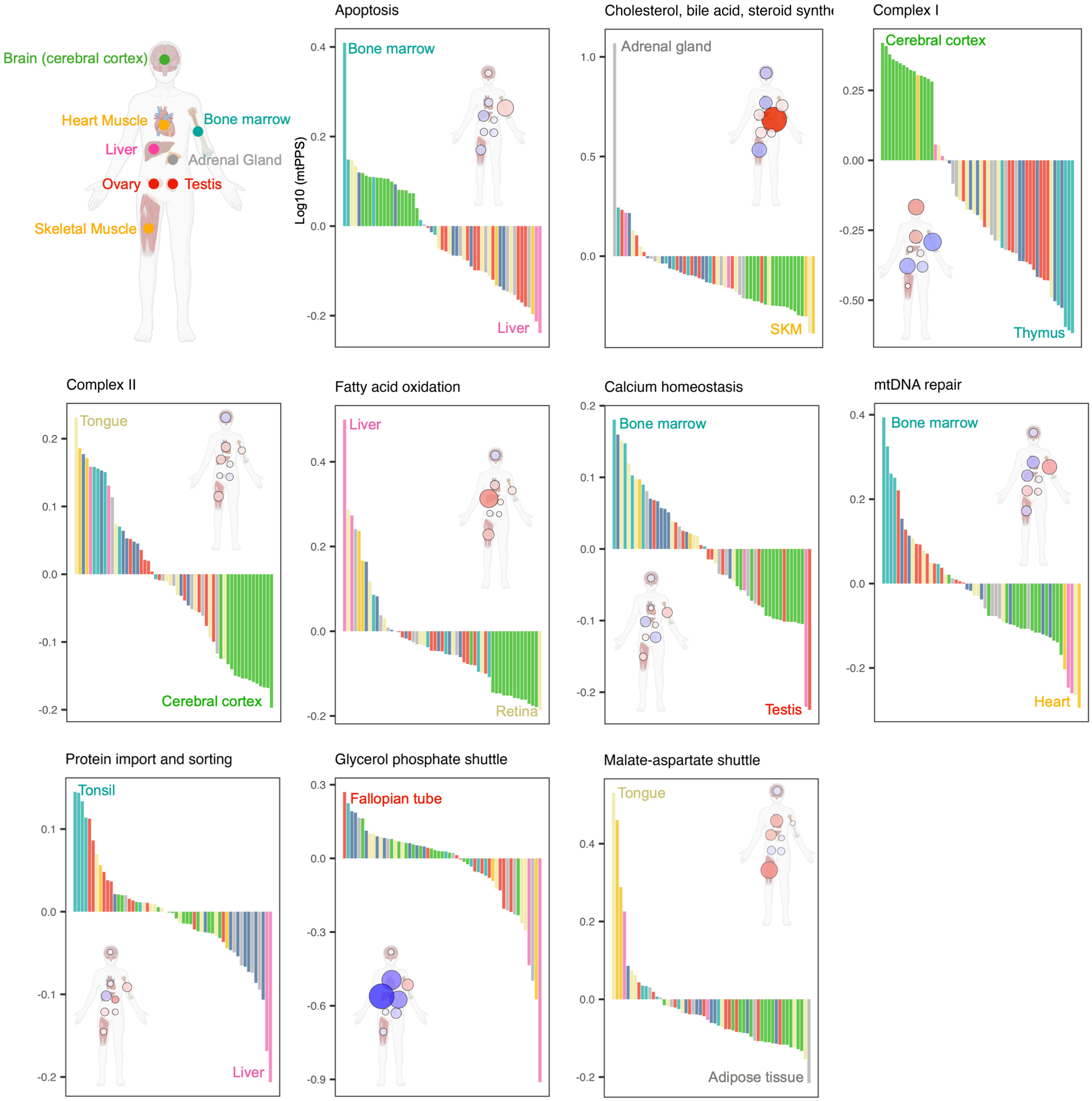

**SUPPL FIGURE S9**

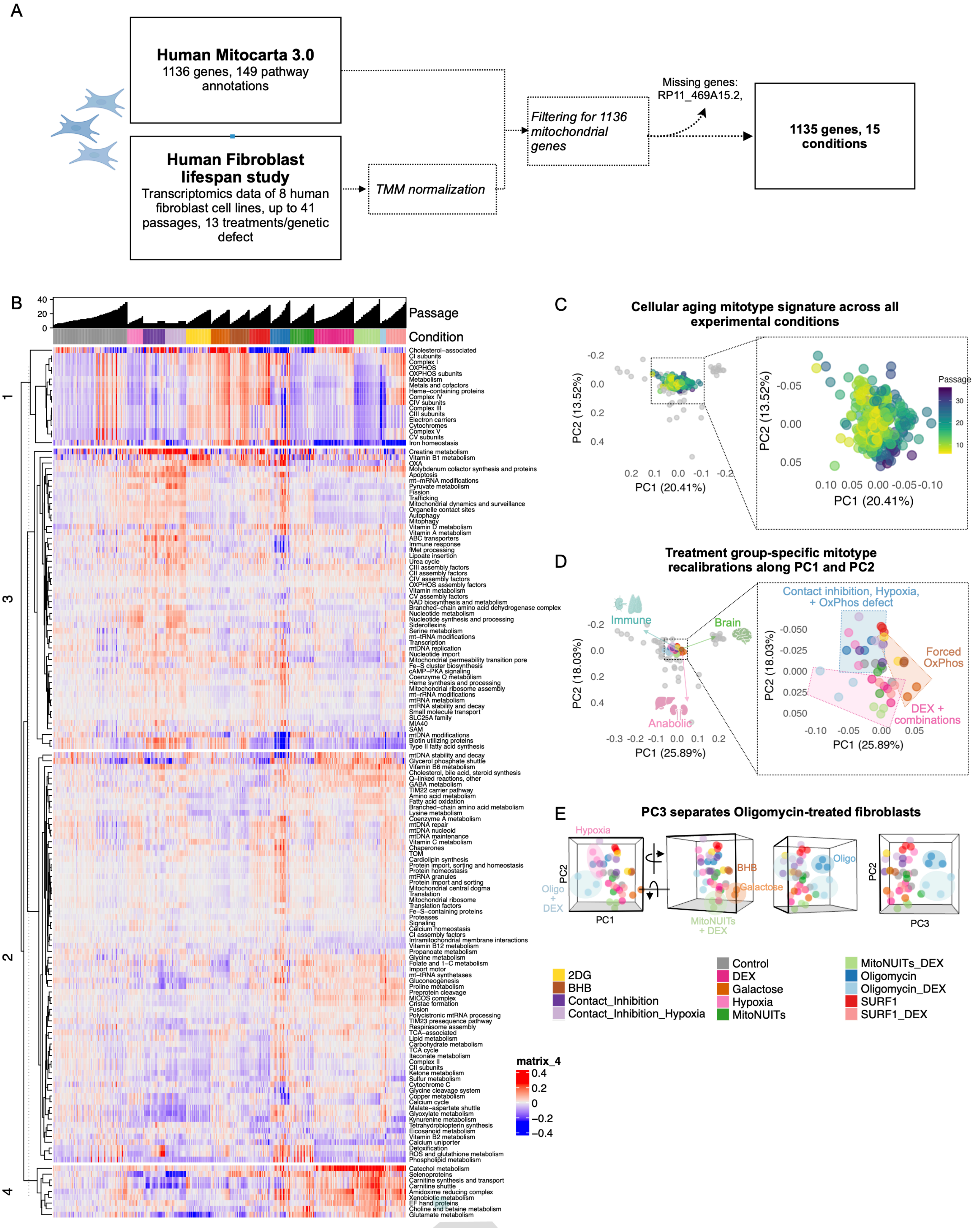

**SUPPL FIGURE S10**

A

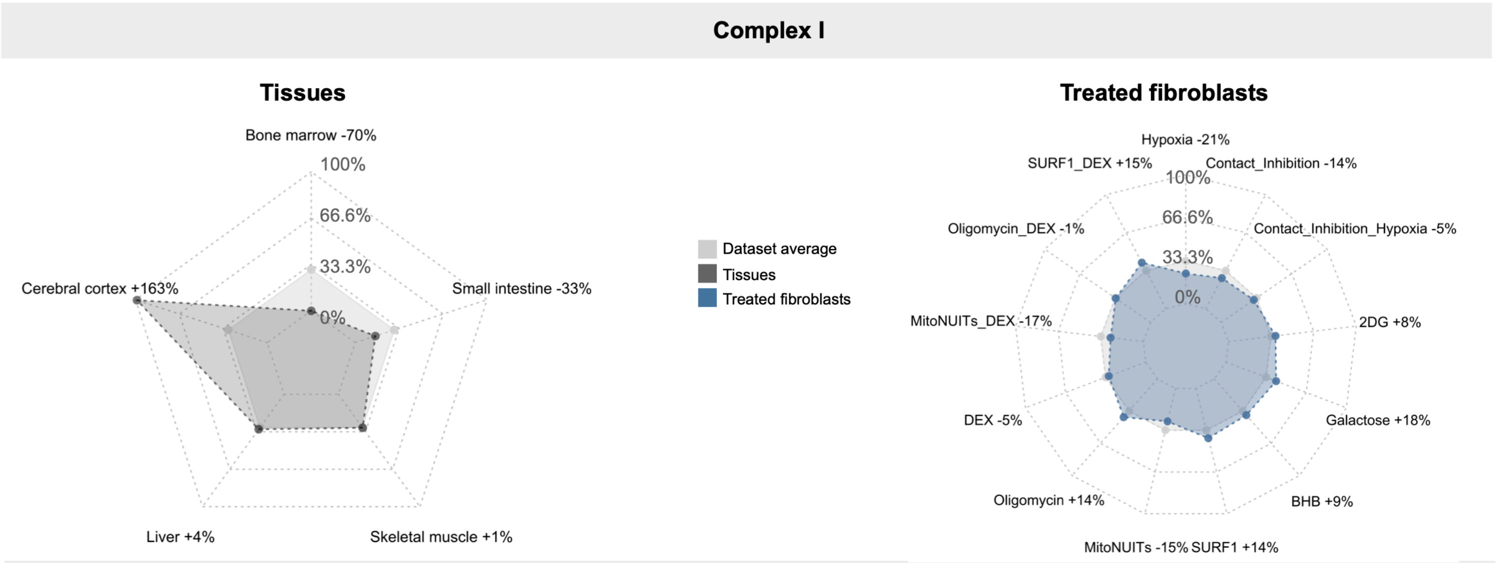


B

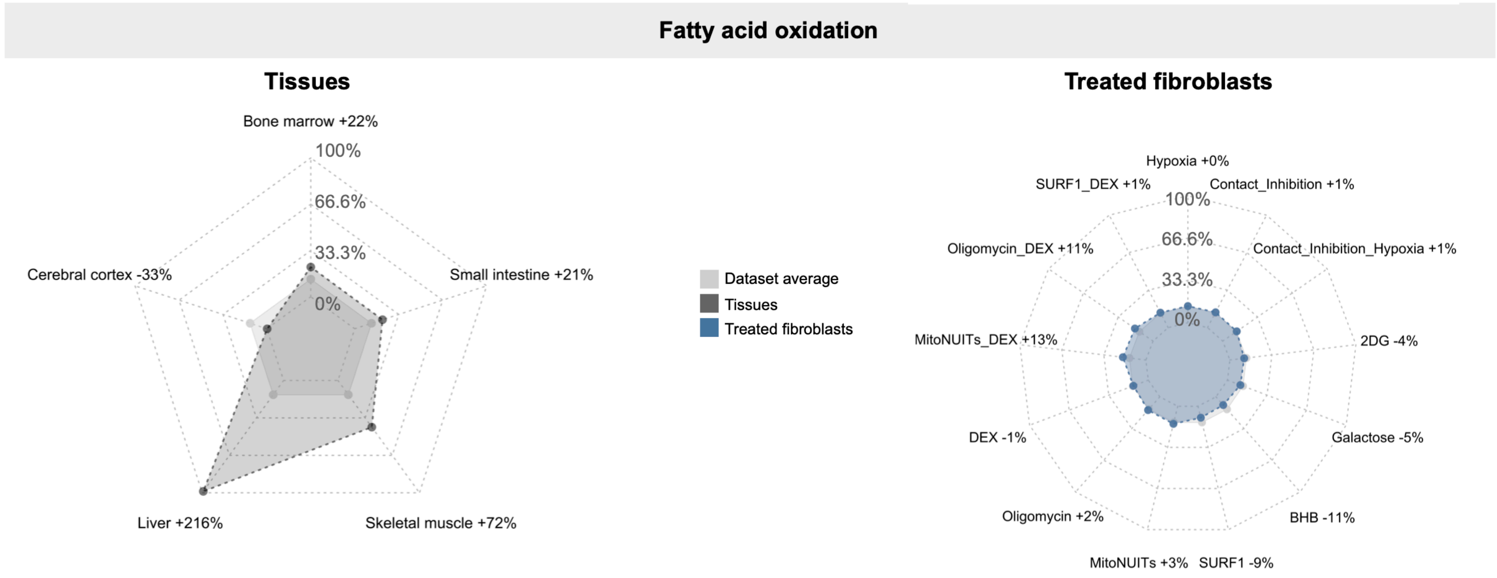


C

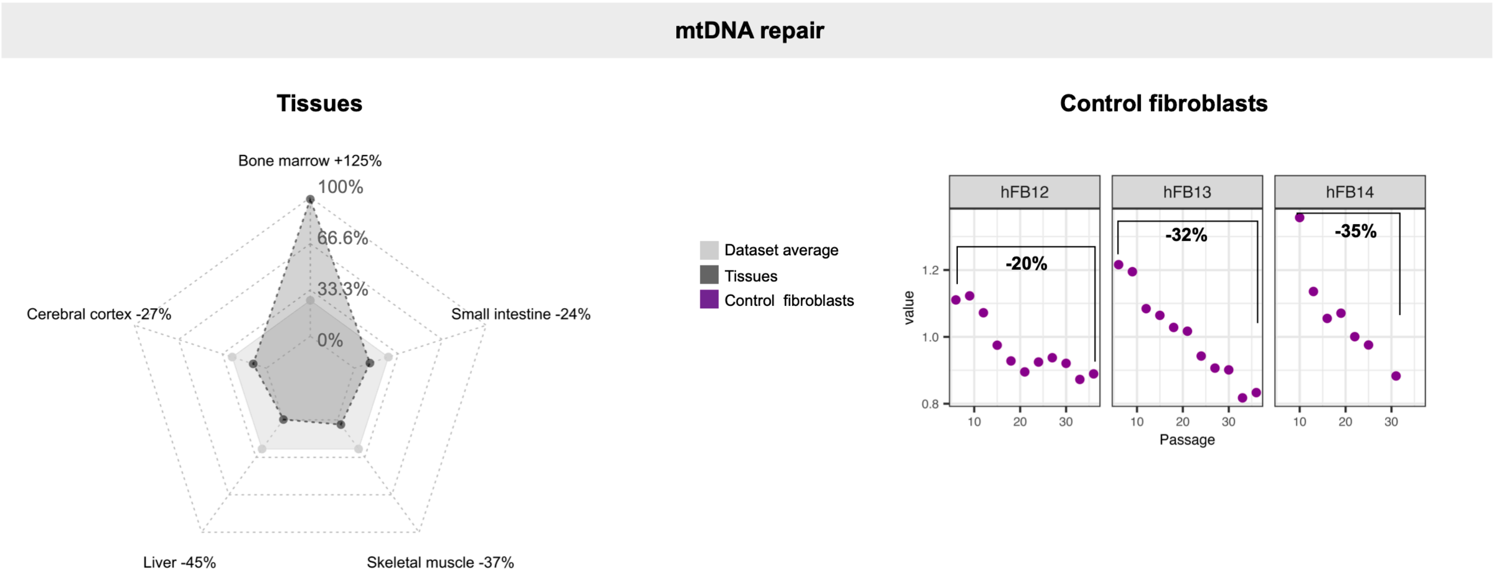

**SUPPL FIGURE S11**

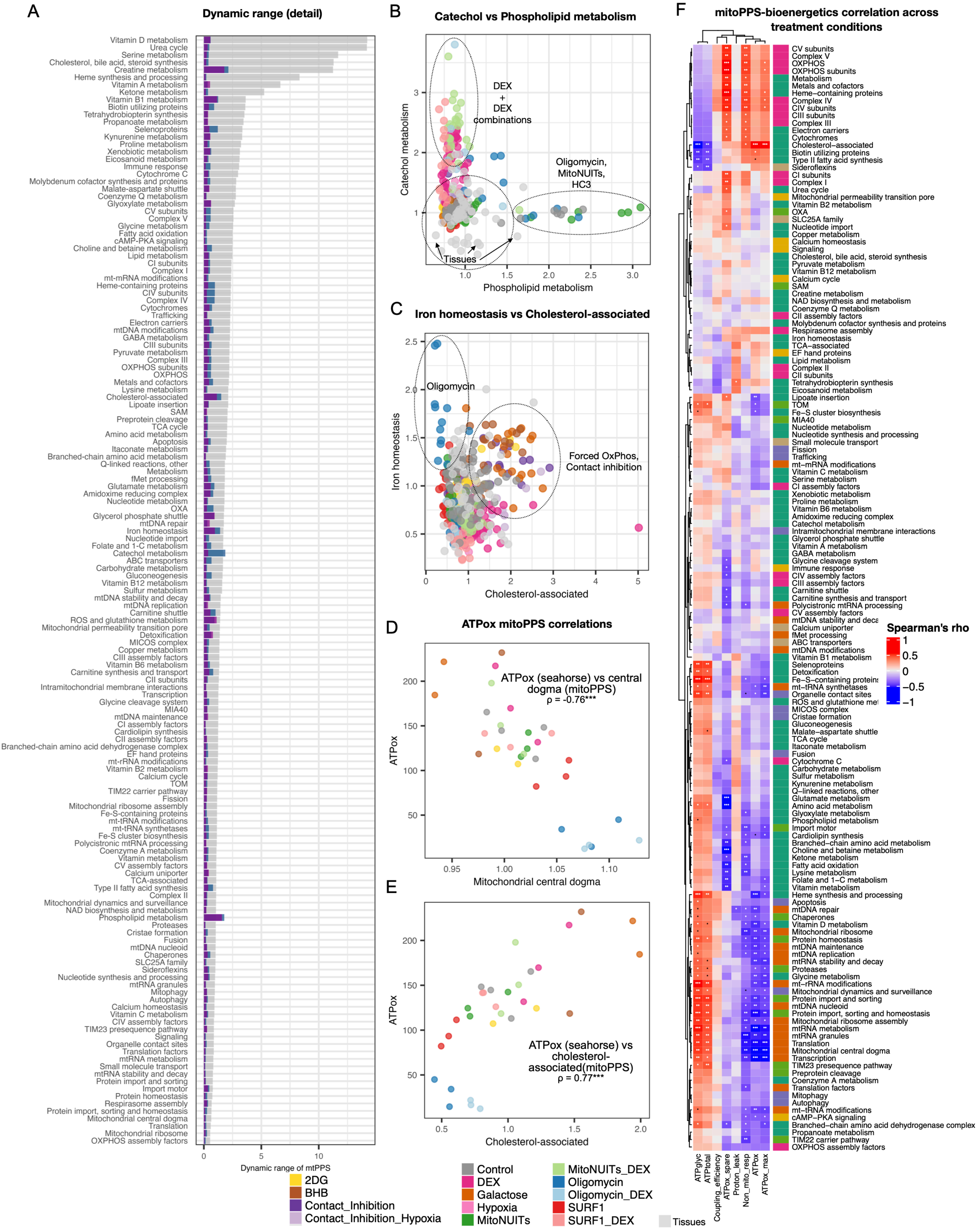

### SUPPL FIGURE S12

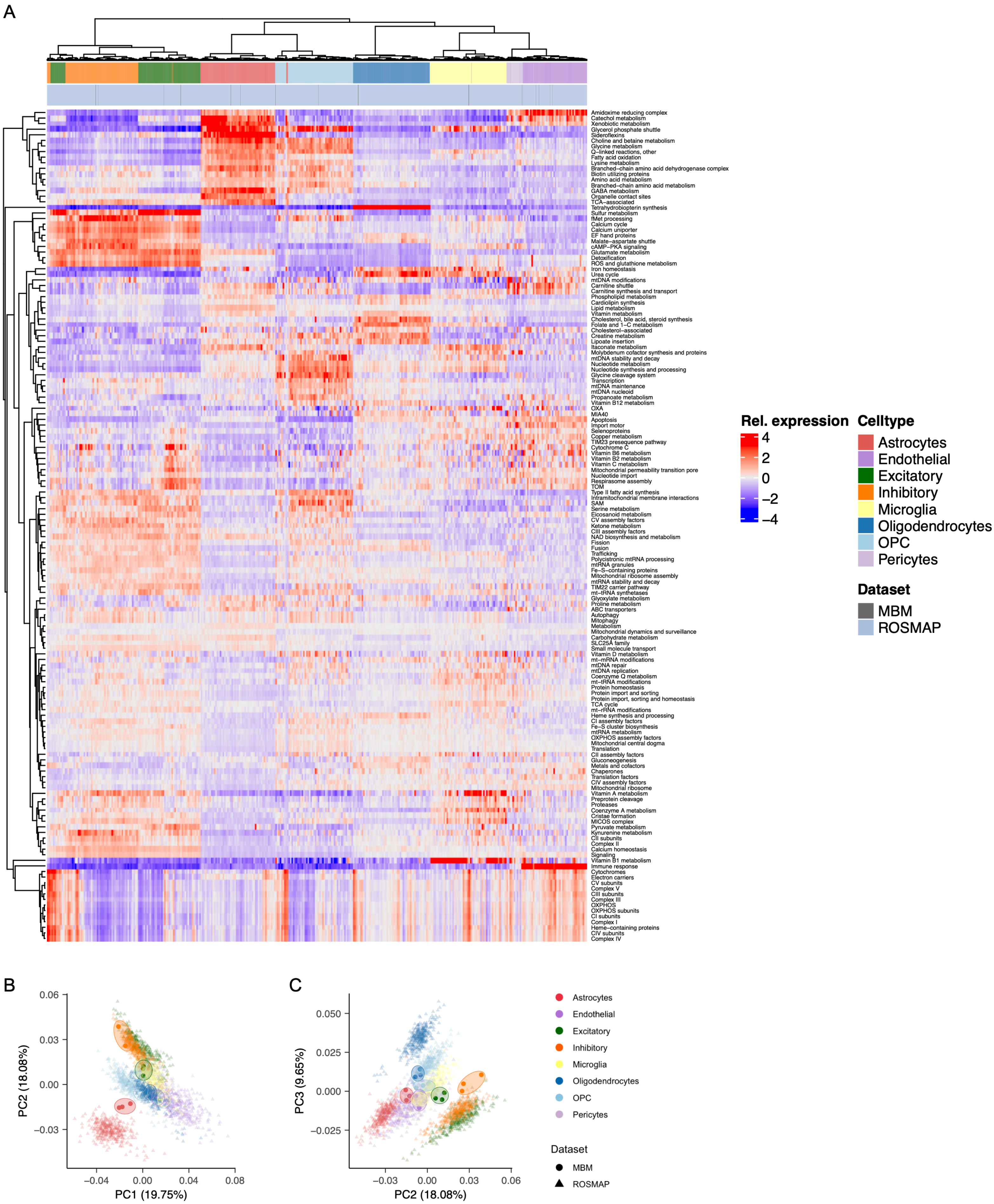

### SUPPL FIGURE S13

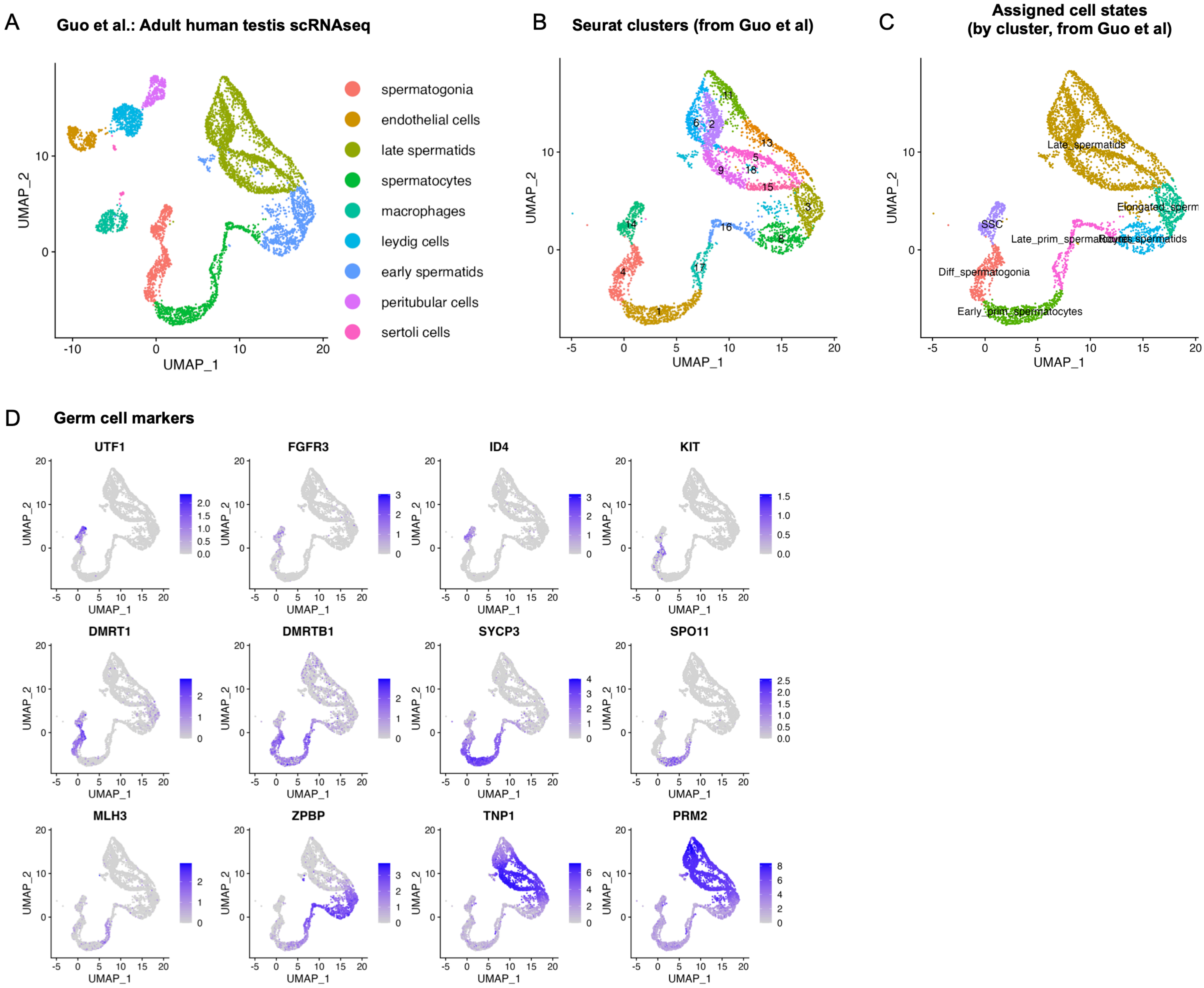

#### SUPPL FIGURE S14

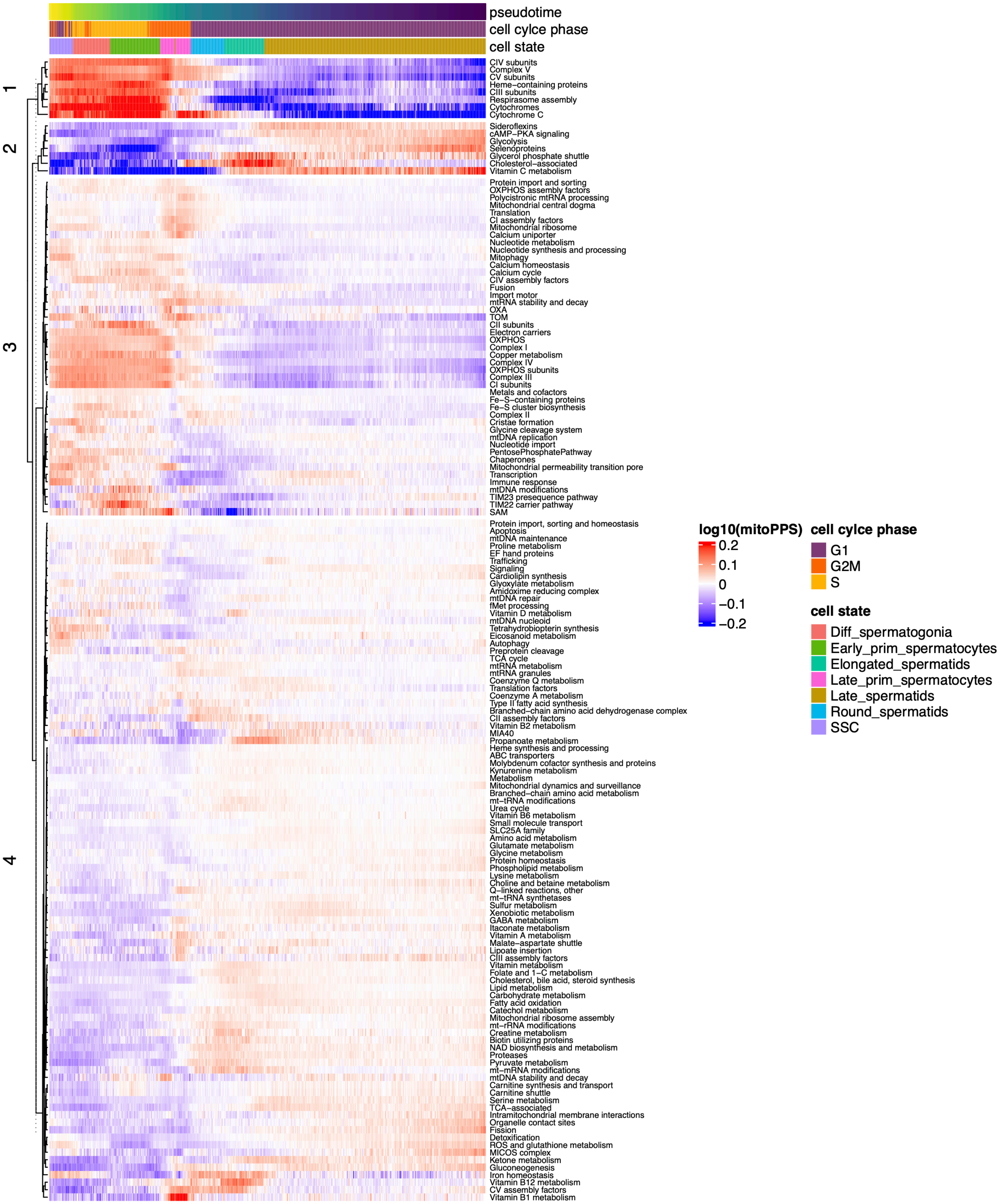

#### SUPPL FIGURE S15

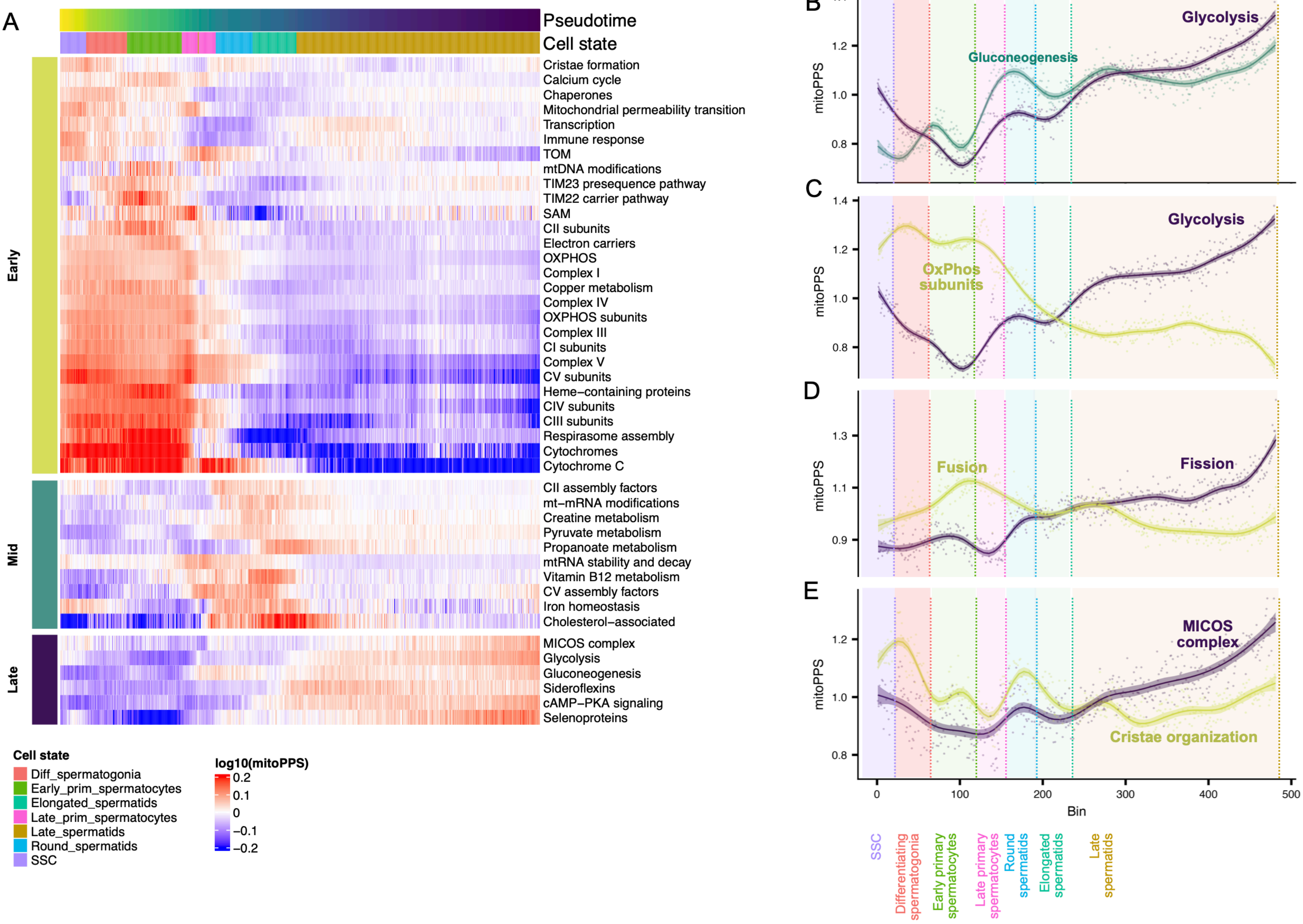
